# Bayesian Modeling Reveals Ultrasensitivity Underlying Metabolic Compensation in the Cyanobacterial Circadian Clock

**DOI:** 10.1101/835280

**Authors:** Lu Hong, Danylo O Lavrentovich, Archana Chavan, Eugene Leypunskiy, Eileen Li, Charles Matthews, Andy LiWang, Michael J Rust, Aaron R Dinner

## Abstract

Mathematical models can enable a predictive understanding of mechanism in cell biology by quantitatively describing complex networks of interactions, but such models are often poorly constrained by available data. Owing to its relative biochemical simplicity, the core circadian oscillator in *Synechococcus elongatus* has become a prototypical system for studying how collective dynamics emerge from molecular interactions. The oscillator consists of only three proteins, KaiA, KaiB, and KaiC, and near-24-h cycles of KaiC phosphorylation can be reconstituted in vitro. Here, we formulate a molecularly-detailed but mechanistically agnostic model of the KaiA-KaiC subsystem and fit it directly to experimental data within a Bayesian parameter estimation framework. Analysis of the fits consistently reveals an ultrasensitive response for KaiC phosphorylation as a function of KaiA concentration, which we confirm experimentally. This ultrasensitivity primarily results from the differential affinity of KaiA for competing nucleotide-bound states of KaiC. We argue that the ultrasensitive stimulus-response relation is critical to metabolic compensation by suppressing premature phosphorylation at nighttime.

**Synopsis:** This study takes a data-driven kinetic modeling approach to characterizing the interaction between KaiA and KaiC in the cyanobacterial circadian oscillator and understanding how the oscillator responds to changes in cellular metabolic conditions.

- An extensive dataset of KaiC autophosphorylation measurements was gathered and fit to a detailed yet mechanistically agnostic kinetic model within a Bayesian parameter estimation framework.
- KaiA concentration tunes the sensitivity of KaiC autophosphorylation and the period of the full oscillator to %ATP.
- The model reveals an ultrasensitive dependence of KaiC phosphorylation on KaiA concentration as a result of differential KaiA binding affinity to ADP- vs. ATP-bound KaiC.
- Ultrasensitivity in KaiC phosphorylation contributes to metabolic compensation by suppressing premature phosphorylation at nighttime.

## Introduction

Achieving a predictive understanding of biological systems and chemical reaction networks is challenging because complex behavior can emerge from even a small number of interacting components. Classic examples include the propagation of action potentials in neurobiology and chemical oscillators such as the Belousov–Zhabotinsky reaction. The collective dynamics in such systems cannot be easily intuited through qualitative reasoning alone, and thus mathematical modeling has long played an important role in summarizing and interpreting existing observations and formulating testable, quantitative hypotheses.

In general, mathematical modeling can be classified as either “forward” or “reverse.” In forward modeling, known interactions are expressed mathematically, which allows a researcher to draw out the logical implications of the model and its underlying assumptions (Gunawardena, 2014). In reverse modeling, experimental data are used to infer unknown interactions through a statistical approach (Villaverde and Banga, 2014). Many forward modeling studies are highly phenomenological; such studies excel in showing how effects like feedback (Novák and Tyson, 2008) and ultrasensitivity (Ferrell and Ha, 2014a,b) can give rise to collective dynamics, including bistable switching, oscillation, and adaptation (Ma et al., 2009). The simplicity of this class of models, however, makes quantitative prediction and experimental verification difficult. Reverse modeling, on the other hand, has found success in untangling complex interactions in -omic data (Machado et al., 2011; Wu et al., 2017) and signaling pathways such as the eukaryotic circadian clock (Forger and Peskin, 2003) and the JAK2/STAT5 signaling pathway (Hug et al., 2013). However, the complexity of such models raises issues of identifiability, i.e., whether a model topology and/or parameter values can be uniquely determined given the input data (Bellman and Åström, 1970; Cobelli and DiStefano, 1980). Furthermore, the nonlinear dynamics typical of such models give rise to non-convex optimization problems that pose significant technical and computational challenges.

The circadian clock from the cyanobacterial species *Synechococcus elongatus* PCC 7942 (Johnson et al., 2011) represents a unique opportunity to combine elements of both forward and reverse modeling. The core oscillator is post-translational (Tomita et al., 2005) and consists of just three proteins: KaiA, KaiB, and KaiC. A stable rhythm in KaiC phosphorylation with a period of nearly 24-h emerges spontaneously from these components, driven by KaiA-dependent autokinase reactions followed by a KaiB-mediated delayed negative feedback loop that favors dephosphorylation. The phosphorylation cycle can be reconstituted in vitro while still retaining the hallmarks of circadian rhythms in living organisms (Nakajima et al., 2005; Yoshida et al., 2009; Rust et al., 2011; Leypunskiy et al., 2017). Previous work has clearly articulated the basic biochemical events in the phosphorylation cycle (Johnson et al., 2011; Swan et al., 2018), allowing specification of a model topology with few ambiguities.

Despite the apparent simplicity of the system, the dynamics of the Kai oscillator are sufficiently complex that reverse modeling can provide useful insights. KaiC molecules can exist in multiple phosphorylation states and nucleotide-bound states, and how these states affect KaiC’s interaction with KaiA (Mori et al., 2018) and KaiB (Phong et al., 2013; Lin et al., 2014) is not fully understood. A related unresolved issue is the effect of the solution nucleotide pool (ATP and ADP) on the oscillator. In *S. elongatus*, the day/night cycle is reflected in the cellular metabolic state, including changes in the adenylate nucleotide pool %ATP (defined as 100%[ATP]/([ATP] + [ADP])), which acts as a timing cue and plays an important role in controlling the amplitude and phase of the phosphorylation cycle (Rust et al., 2011; Phong et al., 2013; Leypunskiy et al., 2017). KaiC is an ATPase (Terauchi et al., 2007) and phosphotransferase (Nishiwaki and Kondo, 2012), and its activities are regulated by which nucleotides are bound. The nucleotide-bound state is in turn regulated by KaiA, which acts as a nucleotide-exchange factor (Nishiwaki-Ohkawa et al., 2014). The kinetics of nucleotide exchange, the affinities of KaiC for nucleotides, and the heterogeneity of nucleotide-bound states in the KaiC hexamer have been measured (Nishiwaki-Ohkawa et al., 2014; Abe et al., 2015), but it is experimentally challenging to monitor all of the relevant quantities simultaneously over the course of the cycle.

Here we take a data-driven Bayesian modeling approach (**Figure 1**A) to elucidate the regulatory relations between KaiA, nucleotides in solution, KaiC phosphorylation, and KaiC nucleotide-bound state, with the goal of deducing dynamical rules that can predict the behavior of the system. The resulting model does not include KaiB; it focuses on describing the dynamics of phosphorylation during the daytime part of the clock cycle. To provide a training set for this model, we collected kinetic time series characterizing the metabolic sensitivity of the KaiC phosphorylation kinetics (in the absence of KaiB) over a wide range of KaiA concentrations ([KaiA]) and %ATP. Although such data do not give us direct access to all relevant states of the KaiA-KaiC subsystem, they place constraints on the underlying molecular interactions. Bayesian parameter estimation (MacKay and Kay, 2003) has been used to systematically quantify parameter uncertainties and compare models in many fields (Geweke, 1989; Wasserman, 2000; Hou et al., 2012), including systems biology (Flaherty et al., 2008; Klinke, 2009; Toni Tina et al., 2009; Xu et al., 2010; Schmidl et al., 2012; Eydgahi et al., 2013; Pullen and Morris, 2014; Mello et al., 2018). Here it allows us to estimate parameter values, quantify the importance of specific model elements, and make mechanistic predictions from the model.

**Figure 1:**
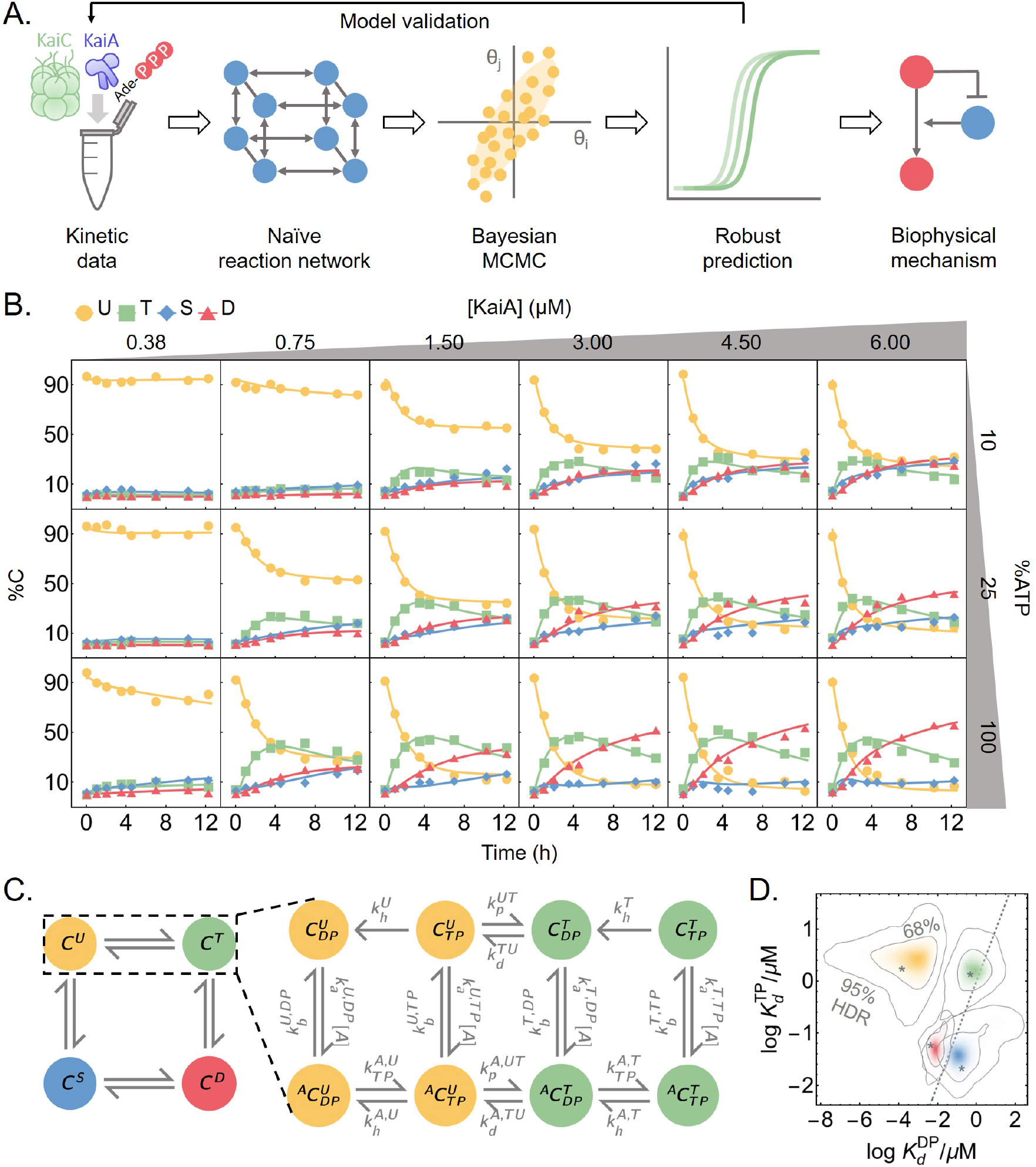
Phosphorylation data are fit by a mechanistically agnostic kinetic model. A) An outline of the data-driven Bayesian model fitting approach employed in this work. B) To constrain the model, measurements of KaiC phosphorylation kinetics were collected at six [KaiA] and three %ATP conditions. The curves represent the best fit model prediction. C) A schematic of the mass action kinetics model. The model elaborates on the autophosphorylation reactions of KaiC by explicitly keeping track of the time evolution of the KaiC phosphoforms, nucleotide-bound states, and KaiA binding mediated by phosphotransfer, nucleotide exchange, and ATP hydrolysis. Note that the KaiA binding reactions are second-order, but KaiA concentration ([A]) is written as part of the effective first-order rate constant. See the main text for a discussion of the state and rate constant nomenclature and **Figure S1**A for a schematic of the full model. The posterior distributions for log KaiA dissociation constants (base 10). The horizontal axis represents the affinity for ADP-bound KaiC, and the vertical axis represents the affinity for ATP-bound KaiC; the four colors correspond to the KaiC phosphoforms, as in panel C. The asterisks represent the best fit, and the contour lines represent the 95% and 68% highest posterior density regions (HDR). The dashed line represents the 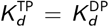 line, so that densities above the line indicate higher affinity for the ADP-bound species and densities below the line indicate higher affinity for the ATP-bound species.

The Markov chain Monte Carlo (MCMC) sampling method that we use to fit the model to the data yields an ensemble of parameter sets, rather than a single best fit. We find that, even with extensive training data, many microscopic parameters in the model are not tightly constrained and their values vary widely across the ensemble of fits. Despite this, we show that this ensemble of fits robustly makes predictions that are borne out in experimental tests (Brown and Sethna, 2003; Gutenkunst et al., 2007). In particular, the model reveals an ultrasensitive dependence of phosphorylation on the concentration of KaiA, with strong nonlinearity at low [KaiA], conditions that likely apply near the nighttime to daytime transition point, when a large fraction of KaiA molecules are inhibited. The ultrasensitive response primarily arises from a differential affinity of KaiA for different nucleotide-bound states of KaiC. This mechanism is analogous to substrate competition (Ferrell and Ha, 2014b), where kinetic competition of multiple enzyme substrates leads to ultrasensitivity.

Lastly, we consider the implications of these results for the full oscillator, in which KaiC rhythmically switches between phosphorylation and dephosphorylation. Incorporation of the ultrasensitive response to KaiA into a mathematical model of the full oscillator suggests that this effect both stabilizes the period against changes in the nucleotide pool and allows oscillations to persist even when KaiB binds KaiA relatively weakly. Consistent with this prediction, we find that a substantial amount of KaiA is not bound by KaiB even when the clock is dephosphorylating. These results shed new light on metabolic compensation, a property that allows robust 24-h oscillation in spite of changes in %ATP conditions (Johnson and Egli, 2014). Taken together, our results show how the Bayesian framework combined with extensive training data can be used to discover unanticipated mechanisms and direct experimental investigations.

## Results

### A molecularly motivated model of KaiA-KaiC dynamics

To probe the response of KaiC phosphorylation to a wide range of metabolic conditions, we made kinetic measurements of KaiC phosphorylation at three %ATP conditions and six [KaiA] conditions while holding the KaiC concentration constant (**Figure 1**B). KaiC is a homohexamer and each subunit has two domains, termed CI and CII. Both CI and CII domains have ATPase activity (Hayashi et al., 2003; Pattanayek et al., 2004; Terauchi et al., 2007), while the CII domain is in addition a bidirectional phosphotransferase (Nishiwaki and Kondo, 2012; Egli et al., 2012) with two phosphorylation sites (Xu et al., 2004; Rust et al., 2007; Nishiwaki et al., 2007). Each KaiC subunit thus has four phosphoforms: the unphosphorylated (U), phosphoserine-431 (S), phosphothreonine-432 (T), and doubly phosphorylated (D) states. The measurement resolves the kinetics of all four KaiC phosphoforms.

Our strategy is to fit these data with a model of the KaiC catalytic cycle with a minimum of simplifying assumptions. To this end, we formulate a model based on mass action kinetics. We explicitly keep track of three properties of the CII domain of each KaiC subunit: its phosphorylation status (right superscripts in **Figure 1**C), nucleotide-bound state (right subscript), and whether or not KaiA is bound (left superscript). We do not consider CI or the hexameric nature of KaiC explicitly (see **SI** for further discussion). There are thus 16 possible KaiC states, 8 of which are shown in **Figure 1**C, along with the phosphotransfer, nucleotide exchange, KaiA (un)binding, and hydrolysis reactions that connect the states (see **Figure S1**A for the full model structure). We also hypothesized that nucleotides might interact directly with KaiA, which could allow KaiA’s activity to directly depend on nucleotides in solution. However, we did not detect any direct interaction between KaiA and ATP or ADP using NMR spectroscopy, so we do not allow for this scenario in the model (**Figure S2**). Below, we step through the four classes of reactions that we include; further details can be found in Materials and Methods.

#### Phosphotransfer

KaiC is a bidirectional phosphotransferase (Egli et al., 2012; Nishiwaki and Kondo, 2012), which means that it can transfer a γ-phosphate group from a bound ATP to a phosphorylation site, but unlike a typical phosphatase, it regenerates ATP from ADP during dephosphorylation, i.e.,

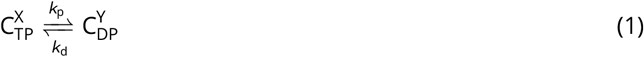

where (X, Y) ∈ {(U, T), (U, S), (T,D), (S, D)}. This mechanism implies that the nucleotide-bound state of KaiC has a significant impact on the direction of its phosphotransferase activity: an ATP-bound KaiC presumably cannot dephosphorylate, and an ADP-bound KaiC cannot phosphorylate.

#### Nucleotide exchange

KaiA binding to the CII domain (Kim et al., 2008; Pattanayek and Egli, 2015) stimulates KaiC autophosphorylation (Iwasaki et al., 2002; Williams et al., 2002; Kageyama et al., 2006). Recent work has shown that KaiA can bind to KaiC and act as a nucleotide-exchange factor (Nishiwaki-Ohkawa et al., 2014) by facilitating conformational changes at the subunit interface that promote solvent exposure of the nucleotide-binding pocket (Hong et al., 2018). It is currently unclear whether this nucleotide exchange activity is responsible for all of KaiA’s effect on KaiC or whether it alters the KaiC catalytic cycle in other ways (see **SI** for further analysis of this issue). The reversible binding of KaiA

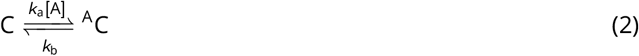

contributes two classes of rate constants, *k*_a_ and *k*_b_.

Because the CII domain of KaiC releases its bound nucleotide very slowly in the absence of KaiA (Nishiwaki-Ohkawa et al., 2014), we ignore the possibility of KaiA-independent nucleotide exchange in the model. Under the assumptions that i) the apo state is in a quasi-steady state, ii) the ADP and ATP on-rates are identical, and iii) ATP release is slow, nucleotide exchange can be modeled as a one-step reaction

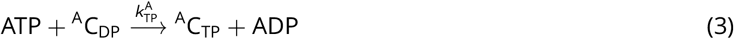

where

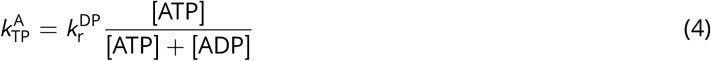

and 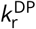 is the ADP dissociation rate constant. Nucleotide exchange thus contributes one class of rate constant, 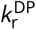. See Materials and Methods for the derivation of (4).

#### ATP hydrolysis

Finally, we allow for irreversible ATP hydrolysis in the CII domain

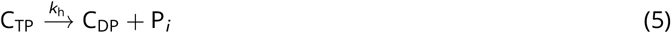

which contributes one class of rate constants, *k*_h_. Because each KaiC molecule consumes relatively little ATP on the timescale of a day (Terauchi et al., 2007), we assume the solution ATP and ADP concentrations are constant.

#### Species-dependent rates

Given the six classes of rate constants, *k*_p_, *k*_d_, *k*_a_, *k*_b_, 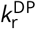, and *k*_h_, we make the model maximally general, or mechanistically agnostic, by allowing each rate constant to potentially depend on the specific molecular state involved in the reaction. For example, the KaiA dissociation rate constant is allowed to vary depending on the nucleotide-bound state and phosphoform background of KaiC, and thus the dissociation rate constants for the ADP-bound U 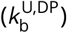 and ATP-bound T phosphoforms 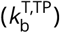 are two independent model parameters. In this way, the parameter fitting and model comparison procedures automatically test specific biochemical hypotheses about the functions of KaiA and KaiC. For example, allowing the KaiA off-rates to depend on the nucleotide-bound states is equivalent to the hypothesis that KaiA has different dwell times for ATP-versus ADP-bound states of KaiC. In fact, because each reaction has an independent rate constant, except for the thermodynamic constraints of detailed balance, the fitting procedure effectively allows for simultaneous testing of all possible two-way interactions of the three categories of KaiC properties, without a priori preference for any particular mechanism.

### The data constrain the parameters to widely varying degrees

We estimate the model parameters through a Bayesian framework. In this framework, we maximize the posterior probability, which is proportional to the product of the prior distribution and the likelihood function. Here, we interpret the prior as representing subjective beliefs on the model parameters before experimental inputs, while the likelihood function quantifies the goodness of fit. Bayesian parameter estimation reduces to least-squares fitting under the assumption of normally distributed residuals and uniform priors. In practice, we find that direct numerical optimization of the posterior usually results in fits that are trapped in low probability local maxima (**Figure S3**B). Thus we instead draw parameters from the prior distribution and then use a heuristic combination of Markov chain Monte Carlo (MCMC) sampling and optimization (Powell’s algorithm) to explore the parameter space. The MCMC method that we use (Goodman and Weare, 2010; Foreman-Mackey et al., 2013) efficiently searches the parameter space by simulating an ensemble of parameter sets in parallel; the spread of the ensemble reflects the geometry of the posterior distribution and is used to guide the directions of Monte Carlo moves. See Materials and Methods for a more mathematical treatment of the fitting procedure and comparison of different numerical optimization and sampling methods.

We use this approach to fit the phosphorylation data (**Figure 1**B) together with previously published data on dephosphorylation (Rust et al., 2011), ATP hydrolysis rate (Terauchi et al., 2007), and the KaiA dwell time for each KaiC phosphoform (Kageyama et al., 2006; Mori et al., 2018) (see Materials and Methods). Overall, the model achieves excellent agreement with the training data (**Figure 1**B and **Figure S4**A–C). In the following analyses, we refer to model predictions using the best fit parameter values, and quantify the uncertainties using the posterior distribution (see **SI** for further discussion on the convergence of the simulation).

We find that certain parameters, such as the hydrolysis rates in the U and T phosphoforms and the KaiA off-rates from the U phosphoform, are tightly constrained, while many others, mainly involving S and D phosphoforms, are less constrained, in the sense that their posterior distributions span multiple orders of magnitude, exhibit multimodality, or cannot be reproduced over multiple independent runs (**Figure S1**B). Some parameters are highly correlated and certain combinations of the parameters are much better constrained than the individual parameters. For example, the posterior distributions for the KaiA binding affinities (**Figure 1**D) appear better constrained than the on/off rates (**Figure S5**B).

Taken together, these results are consistent with the notion that collective fits of multiparameter models are generally “sloppy,” meaning that the sensitivities of different combinations of parameters can range over orders of magnitude with no obvious gaps in the spectrum (Brown and Sethna, 2003; Gutenkunst et al., 2007). As we will see, we can nonetheless make useful predictions using the ensemble of model parameters, because the model behavior is constrained along the stiffest directions of the posterior distribution. By contrast, direct parameter measurements need to be both complete and precise to achieve similar predictive validity (Gutenkunst et al., 2007). We further characterize the structure of the parameter space in **SI** and **Figure S5**.

### KaiC (de)phosphorylation goes through transient kinetic intermediates

Given the model, we can interpret the underlying molecular events in KaiC phosphorylation. Here we consider the phosphorylation kinetics at the standard reaction condition (3.5 μM KaiC, 1.5 μM KaiA, 100% ATP; **Figure 2**A and B, solid curves); we examine the effect of varying [KaiA] and %ATP in the following sections. At the beginning of the phosphorylation reaction, KaiC molecules are predominantly in the ADP-bound U state 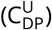, the end product of the dephosphorylation pathway in the absence of KaiA (**Figure 2**A). With the addition of KaiA, the 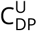 state becomes rapidly depleted within the first 10 minutes of the reaction and enters the 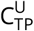 state. Consistent with the kinetic ordering observed in the full oscillator, the 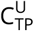 population is primarily converted into the T phosphoform over the S phosphoform. The exact pathway underlying the preference for the T phosphoform is not well constrained by the data, but it appears to be the result of more than just a difference in the relative U→T and U→S phosphorylation rates; there is an interplay between KaiC phosphorylation and KaiA (un)binding kinetics (see **SI** and **Figure S6**). The ADP- and KaiA-bound T phosphoform species are unstable kinetic intermediates, and the population accumulates at the 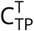 bottleneck for the first 4 hours. As phosphorylation reaches completion, the T phosphoform is converted first into 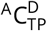 through the unstable ADP-bound intermediates, and then to the 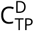 state; the populations of the 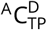 and 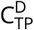 states are comparable at steady state. We note here, however, that previous measurements indicate that approximately 30% of CII nucleotide-binding pockets should be ADP-bound in the presence of KaiA at steady state (Nishiwaki-Ohkawa et al., 2014), which suggests that the stability of the ADP-bound species is systematically underestimated by the model fit.

**Figure 2:**
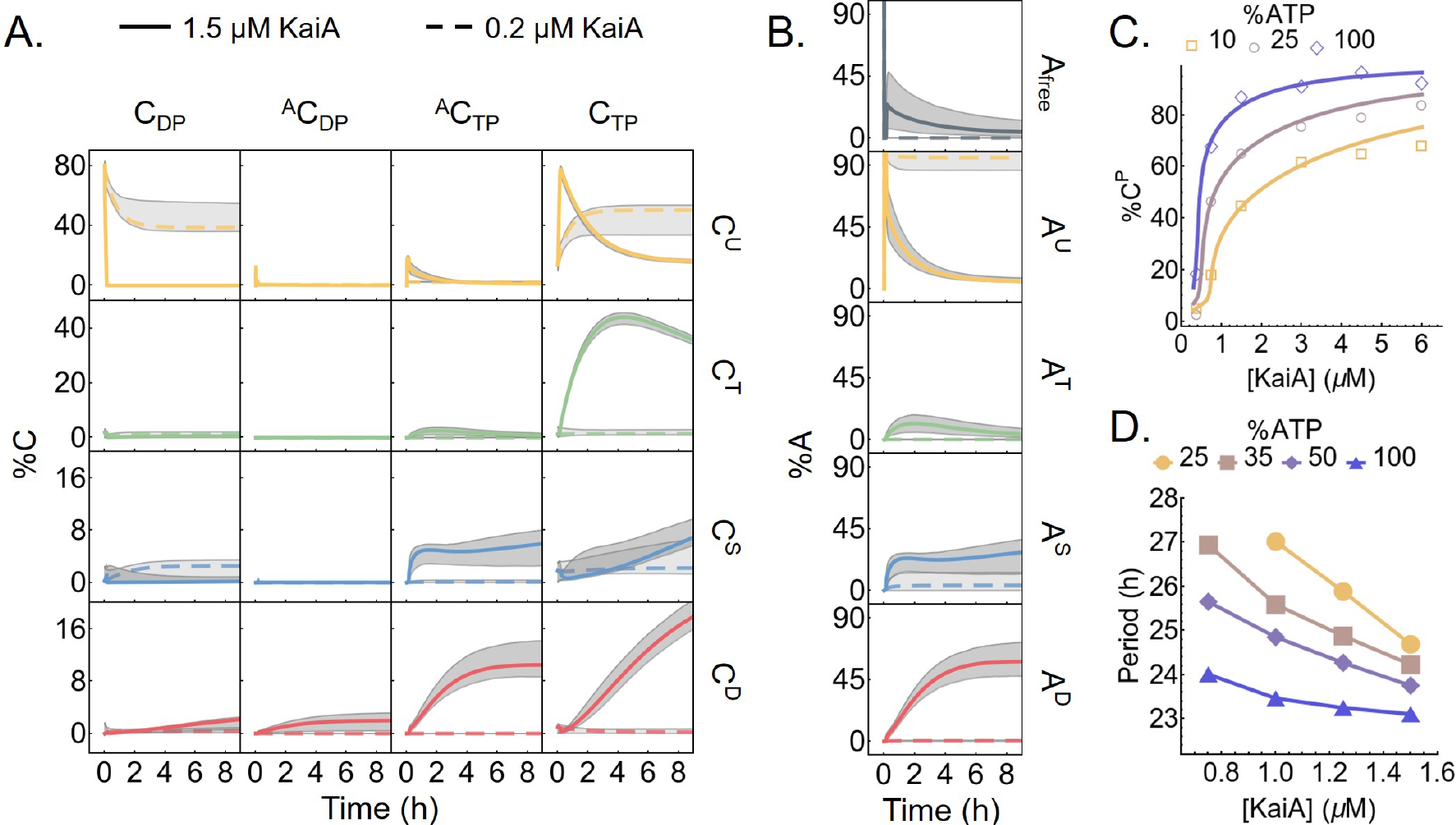
The model captures the kinetics of KaiC phosphorylation. A) The time evolution of all 16 KaiC species in a phosphorylation reaction with 100% ATP and either 1.5 μM (solid curves) or 0.2 μM (dashed curves) KaiA. The corresponding KaiA kinetics, broken down according to the phosphoform of the bound KaiC, is shown in B). The gray regions represent the 95% posterior interval. Refer to **Figure S1**A for the KaiC state nomenclature. C) KaiA concentration can tune the sensitivity of the KaiC phosphorylation level to %ATP. The points represent the measured total percentage phosphorylation levels at *t* = 12:25 h (see **Figure 1**B for the full kinetics), and the curves represent the model prediction at the same time point. D) KaiA concentration can tune the sensitivity of the clock period to %ATP. The period of the full KaiABC oscillator is calculated from fluorescence polarization measurement (see **Figure S7**A-C for further analysis).

During the phosphorylation reaction, the amount of free KaiA is initially transiently depleted due to association with the ADP-bound U phosphoform (**Figure 2**B). Afterwards, KaiA primarily associates with the ATP-bound S and D phosphoforms as they appear, but does not bind to the T phosphoform strongly. Therefore, despite the lack of KaiB in the model, not all KaiA is free during the phosphorylation phase, and the amount of free KaiA depends on both the affinities of the nucleotide-bound states and the mixture of KaiC phosphorylation states (**Figure 1**D).

We were surprised to see that the model fit predicts that the KaiA binding affinity for the ATP-bound T phosphoform is lower than those for the S and D phosphoforms. This is apparently in contradiction with experimental results that show that S-phosphomimetic mutants reduce A-loop exposure and weaken KaiA binding, while T-phosphomimetic mutants have opposite effects (Tseng et al., 2014; Chang et al., 2011). However, there is some ambiguity in the experimental literature, with various results employing different experimental methods and using proteins from different organisms, that has yet to be resolved. Our model stresses the importance of the nucleotide-bound state, especially that of the U phosphoform. KaiC nucleotide-bound state has generally not been measured in KaiA interaction studies and may be different in phosphomimetic mutants (see also Kageyama et al., 2006; Qin et al., 2010a; Murakami et al., 2016).

The dephosphorylation pathway is simpler because KaiA is not involved. In the model, KaiC by itself has no nucleotide exchange activity, and thus phosphorylated KaiC molecules in the absence of KaiA enter a cycle of dephosphorylation by the transfer of phosphoryl groups from the phosphorylation sites back to bound ADP molecules, followed by ATP hydrolysis and removal of inorganic phosphate, until the protein reaches the 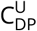 state (**Figure S4**D). The ADP-bound forms of the T, S, and D phosphoforms are only transiently populated, suggesting that the dephosphorylation bottlenecks are the ATP hydrolysis reactions, which make bound ADP available as a cofactor for dephosphorylation, rather than the phosphotransfer reaction itself. The kinetic preference for the D→S dephosphorylation pathway is the direct result of faster dephosphorylation via the D→S compared to the D→T reaction (**Figure S1**B; compare the posterior distribution of 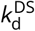 with that of 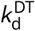). During this process, KaiC can occasionally autophosphorylate, but it is driven irreversibly towards the dephosphorylated state by ATP hydrolysis. We note here that the independence of the dephosphorylation reaction from solution ADP (Rust et al., 2011) is a built-in feature of the model, since solution %ATP only affects the nucleotide exchange rate, which is assumed in the model to be zero in the absence of KaiA.

### KaiA concentration tunes clock sensitivity to %ATP

The model further allows us to summarize and interpret the effect of %ATP and KaiA on KaiC phosphorylation observed in the training dataset (**Figure 2**C). Consistent with previous measurements (Rust et al., 2011; Phong et al., 2013), these results indicate that the near-steady-state (*t* = 12 h) total phosphorylation level of KaiC (%C^P^ = %C^T^ + %C^S^ + %C^D^) is lower in the presence of ADP. Since we simultaneously vary %ATP and [KaiA], the data reveal that this inhibitory effect can be tuned by [KaiA]. In particular, the system is most insensitive to %ATP at either very low or very high [KaiA], while the %ATP sensitivity is the highest around [KaiA] = 0.75 μM.

Interestingly, the %ATP sensitivity of the system cannot be fully abolished even at saturating KaiA concentrations (**Figure 2**C). This effect can be interpreted qualitatively in terms of the structure of the model. When there is more KaiA in solution, more KaiC goes through intermediate states that are in complex with KaiA, by Le Châtelier’s principle. This shifts a larger fraction of the KaiC population through states that allow for exchange of bound ADP for ATP, which promotes phosphorylation. On the other hand, as the %ATP decreases, the ATP to ADP exchange rate decreases according to (4). When nucleotide exchange becomes less efficient, more KaiC stays in ADP-bound states, which are prone to dephosphorylation. In summary, [KaiA] and %ATP both act on the phosphorylation kinetics via the nucleotide exchange step, where %ATP directly regulates the exchange rate constant and sets its upper bound, while [KaiA] controls the population of exchange-competent KaiC and thus the effective exchange rate. Therefore, the effects of KaiA and increasing solution %ATP are not equivalent; because the effective exchange rate cannot exceed the limit set by %ATP, even a saturating amount of KaiA cannot fully compensate for low %ATP.

Given that the metabolic sensitivity of the KaiA-KaiC subsystem can be tuned by KaiA concentration, we asked whether metabolic sensitivity of the full oscillator period may also be tuned by KaiA. To address this question, we characterized the dependence of the period of the in vitro KaiABC oscillator on [KaiA] and %ATP using an optical assay (Leypunskiy et al., 2017; Heisler et al., 2019) that allows automated, parallelized monitoring of the fluorescence polarization of labeled KaiB (**Figure 2**D and **Figure S7**A-C). Consistent with the hypothesis, we found that low KaiA concentration enhances the period sensitivity to %ATP compared to the standard condition (1.5 μM KaiA). These results suggest that the KaiA activity, and how it is controlled, plays a critical role in determining the clock period stability.

### KaiC phosphorylation exhibits ultrasensitive dependence on KaiA levels

In addition to inferring kinetics of states not easily accessible to experiments, the model allows us to interpolate between the training data points and study the relation between KaiC phosphorylation, [KaiA], and %ATP at a much finer resolution. This analysis shows an ultrasensitive dependence of the steady-state %C^P^ on KaiA concentration (**Figure 3**A and D left). Specifically, we see a threshold-hyperbolic stimulus-response relation (Gomez-Uribe et al., 2007; Ferrell and Ha, 2014a), where KaiC phosphorylation is highly suppressed near the sub-micromolar [KaiA] regime, but then follows a right-shifted hyperbolic stimulus-response function once [KaiA] exceeds a threshold. Importantly, the threshold depends on %ATP. The model makes similar predictions for the steady-state T, S, and D phoshoforms as well (**Figure S8**A). However, because of the T→D and S→D phosphotransfer reactions, the stimulus-response relations of T and S are not monotonic functions of [KaiA] because high [KaiA] and high %ATP conditions stabilize the D phosphoform at the expense of the T and S phosphoforms.

**Figure 3:**
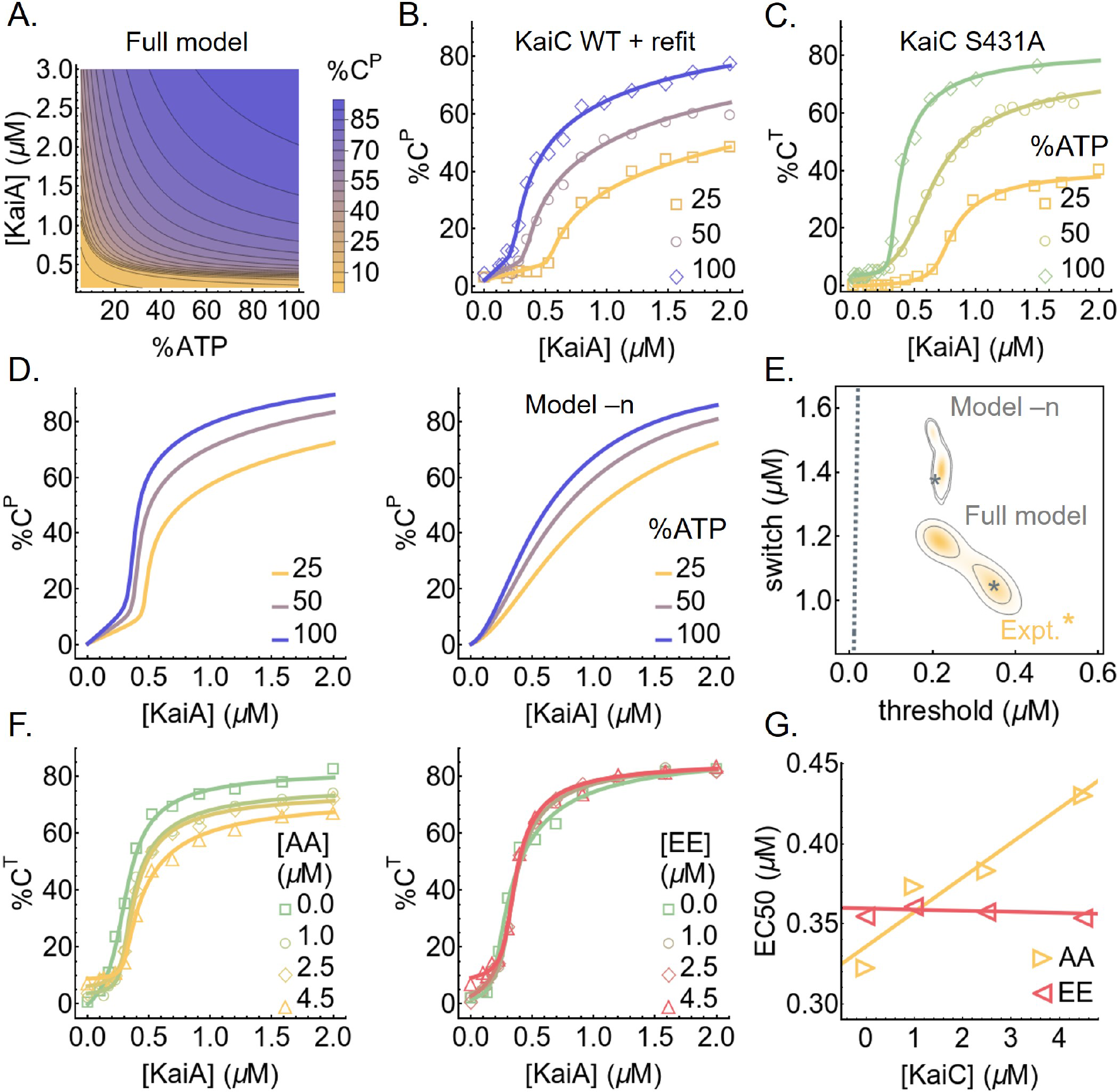
Substrate competition explains KaiC phosphorylation ultrasensitivity. A) The predicted stimulus-response relation of the total steady-state KaiC phosphorylation level as a function of %ATP and [KaiA]. B) Experimentally-determined stimulus-response function of KaiC at three %ATP conditions; the curves are based on refitting the best fit of the full model to the steady-state measurements. C) Similar to B) but for KaiC S431A, which has only one phosphorylation site; the curves are based on independent fits to a simple phenomenological substrate competition model. D) Cross sections of the stimulus-response relation at three %ATP, computed using the full model (left) and model –n (right). E) Posterior distributions for the shapes of the stimulus-response functions at 25% ATP predicted by the full model and model – n. The contours represent the 68% and 95% HDRs, and the gray stars represent the model best fits. The shape of the stimulus-response functions are quantified using two metrics: EC10, which quantifies threshold-like behavior, and EC90 – EC10, which quantifies switch-like behavior. The shape of the experimentally-determined stimulus-response function at 25% ATP is shown as the yellow star. The dashed line represents (EC10, EC90 − EC10) = (*K*/9, 80*K*/9), which characterizes the shape of a hyperbolic stimulus-response function [A]*=*(*K* + [A]) that has no switching or thresholding. F) The stimulus-response functions of KaiC S431A at 100% ATP in the presence of KaiC S431A/T432A (AA; left) and S431E/T432E (EE; right) phosphomimetic mutants to probe the effect of kinetic competition on KaiC phosphorylation. G) The relations between EC50 (the midpoint of a stimulus-response function) and KaiC AA/EE concentrations, quantified using the curves shown in F).

Previous studies of KaiA-KaiC interactions examined the response of KaiC at relatively high KaiA concentrations (≥ 1.2 μM), comparable to the total amount of KaiA used in an oscillating reaction. Ma and Ranganathan (2012) investigated the steady-state stimulus-response relation, but did not consider the effect of %ATP or fully characterize the low [KaiA] regime. Previous reports of initial phosphorylation rates suggest that they exhibit a hyperbolic dependence on [KaiA] (Rust et al., 2007; Lin et al., 2014), similar to simple Michaelis-Menten enzyme-substrate systems. However, this does not imply that the steady-state stimulus-response relation is hyperbolic as well.

To assess the robustness of the model prediction across the ensemble, we use two metrics proposed by Gunawardena (2005) to quantify the shape of the predicted stimulus-response curves for %C^P^ at any fixed %ATP: we use EC10 to measure the extent to which the curve acts as a threshold and EC90 – EC10 to measure the extent to which the curve acts as a switch. Here, EC*x* is the KaiA concentration required to reach *x*% of the steady-state phosphorylation level at saturation. **Figure 3**E shows the distribution of these quantities in the ensemble at 25% ATP. Overall these statistics are tightly constrained by the training data set, and are clearly distinct from those from hyperbolic stimulus-response relations (**Figure 3**E, dashed gray line).

Given the robustness of the prediction, we sought to experimentally verify the shape of the stimulus-response function. We measured KaiC phosphorylation at *t* = 24 h at various concentrations of [KaiA] at three %ATP conditions (**Figure 3**B and **Figure S8**B). Consistent with model prediction, the experimentally-derived stimulus-response relations are ultrasensitive with an %ATP-dependent phosphorylation threshold, and the stimulus-response relation of the S phosphoform at 100% ATP is non-monotonic. We then quantified the shape of the stimulus-response curve for %C^P^ at 25% ATP using the same two metrics defined above (**Figure 3**E, yellow star). At 25% ATP, the shape of the experimentally-derived stimulus-response curve is close to that of the model prediction, but the model fit is systematically less threshold-like (i.e., smaller EC10) and less switch-like (i.e., larger EC90 – EC10). This inconsistency is likely due to a combination of training data under-determining the shape of the curve at the sub-micromolar range (compare **Figure 2**C with **Figure 3**B) and the fitting method under-estimating uncertainties (see **SI**).

Lastly, the saturation phosphorylation levels in the steady-state measurements appear systematically lower than those implied by the training data set (compare **Figure 3**B with D left). This may be a result of batch-to-batch variations in protein and nucleotide quantification. This difference can be corrected by refitting the full model to the steady-state measurement (**Figure 3**B and **Figure S8**C). The refit results suggest that errors in protein and nucleotide concentrations primarily affect the kinetic properties of the S phosphoform in the model (**Figure S8**D), but the refit does not change the qualitative conclusions.

### A substrate competition mechanism underlies ultrasensitivity in KaiC phosphorylation

What is the mechanism of ultrasensitivity in KaiC phosphorylation? Given that each KaiC subunit has two phosphorylation sites, a plausible explanation is multisite phosphorylation, whereby the concentration of the maximally phosphorylated species exhibits an ultrasensitive dependence on the kinase concentration (Gunawardena, 2005) (or in this case, the nucleotide-exchange factor concentration), even if each consecutive phosphorylation step follows mass action kinetics. To examine this possibility, we measured the stimulus-response relation of the KaiC S431A mutant, which has only one phosphorylation site, and the results show ultrasensitivity comparable to that of the WT protein (**Figure 3**C). Furthermore, because KaiC is its own phosphatase, it violates the assumption of distributivity (i.e., at most one modification takes place before the dissociation of the enzyme and substrate) (Gunawardena, 2005). Multisite phosphorylation thus cannot explain the observed ultrasensitivity.

In the ensemble of parameter sets, the KaiA dissociation constant of the ADP-bound (but not ATP-bound) U phosphoform 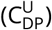 is in or below the nanomolar range, much smaller than that of any other species of KaiC (**Figure 1**D). This is consistent with recent single molecule observations suggesting that the unphosphorylated form of KaiC can bind very tightly to KaiA (Mori et al., 2018) and native mass spectrometry measurements suggesting that KaiA binding to KaiC is enhanced by ATP hydrolysis, which would be needed to produce ADP-bound KaiC (Yunoki et al., 2019). Here, we argue that the key to understanding the origin of ultrasensitivity in the model lies in the differential binding affinity of KaiA to the ADP- and ATP-bound states of KaiC. We note here that since the model does not consider the hexameric structure of KaiC, we cannot rule out possible hexameric cooperative effects that may contribute to ultrasensitivity.

In the model, the differential KaiA binding affinity leads to the following dynamics: KaiA promotes phosphorylation by catalyzing the exchange of the bound ADP for ATP, but this process is in a kinetic competition with ATP hydrolysis, which returns KaiC to the ADP-bound state. At the beginning of the phosphorylation reaction, almost all the KaiA is bound to 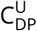 (**Figure 2**A and B) due to its high abundance and high affinity for KaiA (**Figure 1**D). When [KaiA] is low, the competition between nucleotide exchange and hydrolysis in the U phosphoform reaches a steady-state where 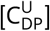 stays above [KaiA] (**Figure 2**A, dashed curves). Therefore, KaiA stays trapped by 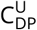 and the phosphorylation products (mostly T) cannot undergo nucleotide exchange. In the absence of KaiA, the autophosphatase activity of KaiC dominates, and the phosphorylation products revert back to the U phosphoform.

When [KaiA] is high, however, the competition between nucleotide exchange and hydrolysis in the U phosphoform pushes 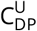 below [KaiA] (**Figure 2**A, solid curves), which frees KaiA to catalyze the nucleotide exchange reactions of the phosphorylation products. Once the flux of phosphorylation, KaiA binding, and nucleotide exchange outweighs that of hydrolysis, dephosphorylation, and KaiA unbinding, the phosphorylation products stay phosphorylated at steady state. Furthermore, the formation of phosphorylation product positively feeds back to deplete 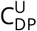, further removing a KaiC state that traps KaiA and leading to rapid saturation of phosphorylation past the [KaiA] threshold. The [KaiA] threshold for phosphorylation depends on %ATP (**Figure 3**A), because when %ATP is low, more KaiA is needed to counteract the reduced ADP-to-ATP exchange rate.

This mechanism is a form of substrate competition (Ferrell and Ha, 2014b; Buchler and Louis, 2008), where the kinetic competition of multiple substrates for enzyme binding leads to ultrasensitivity. Here, KaiA acts as the enzyme, while the ADP-bound U phosphoform and the T (as well as S and D to a lesser extent due to phosphorylation ordering) phosphoform are the substrates that compete for KaiA binding. However, the fact that the phosphorylated and unphosphorylated forms of KaiC can interconvert through phosphotransfer reactions distinguishes the Kai system from a typical substrate competition scheme, where the substrates cannot interconvert.

The model suggests that the U phosphoform plays a special role in generating ultrasensitivity due to the significant difference in the affinity of KaiA for the ATP- vs. ADP-bound states of KaiC (**Figure 1**D). This observation leads to two testable predictions. First, the amount of KaiA required to activate phosphorylation should be higher when more U phosphoform is present. We tested this prediction experimentally by measuring the stimulus-response relation of KaiC S431A in the presence of KaiC S431A/T432A (AA), which mimics the U phosphoform, or KaiC S431E/T432E (EE), which mimics the D phosphoform. The KaiC AA and EE mutants act as competitors for the KaiA-KaiC interaction (**Figure 3**F). Consistent with the hypothesis, the EC50 (i.e., the midpoint of the ultrasensitive switch) is positively correlated with the concentration of KaiC AA, while varying KaiC EE has little effect (**Figure 3**G).

Second, the substrate competition mechanism suggests that the model should exhibit weaker nonlinearity if KaiA has the same affinity to ATP- vs. ADP-bound states of a given KaiC phosphoform. To test this prediction, we constructed simplified models where KaiA on/off rates are set to be independent of the nucleotide-bound state (model –n) or phosphorylation state (model –p) and fit the new models to the experimental data ab initio. Consistent with the prediction, decoupling KaiA on/off rates from the nucleotide-bound states results in a significant loss of ultrasensitivity (**Figure 3**D right and **Figure S9**A). Model –p by contrast behaves similarly to the full model (**Figure S9**C); consistent with the substrate competition mechanism, the ADP-bound states of KaiC in model –p have higher affinity to KaiA than the ATP-bound states, regardless of the phosphorylation state (**Figure S9**C). We quantify the effects of such model reductions by computing the Bayes factor, which is a metric for systematic model comparison that favors goodness of fit but penalizes model complexity and parameter fine tuning (MacKay and Kay, 2003); it is similar to the Bayesian information criterion (Schwarz, 1978), but makes no asymptotic assumptions. The analysis shows that the loss of ultrasensitivity in model –n degrades the fit quality significantly, while model –p is only marginally worse than the full model (**Table 1**). Interestingly, a model where the KaiA on/off rates are completely independent of the state of KaiC (model –n,–p; **Figure S9**D and E) is much worse than either model –n or model –p (**Table 1**). We conclude that the nucleotide-bound state of KaiC plays a key role in regulating its interaction with KaiA and thus in determining phosphorylation kinetics.

**Table 1:**
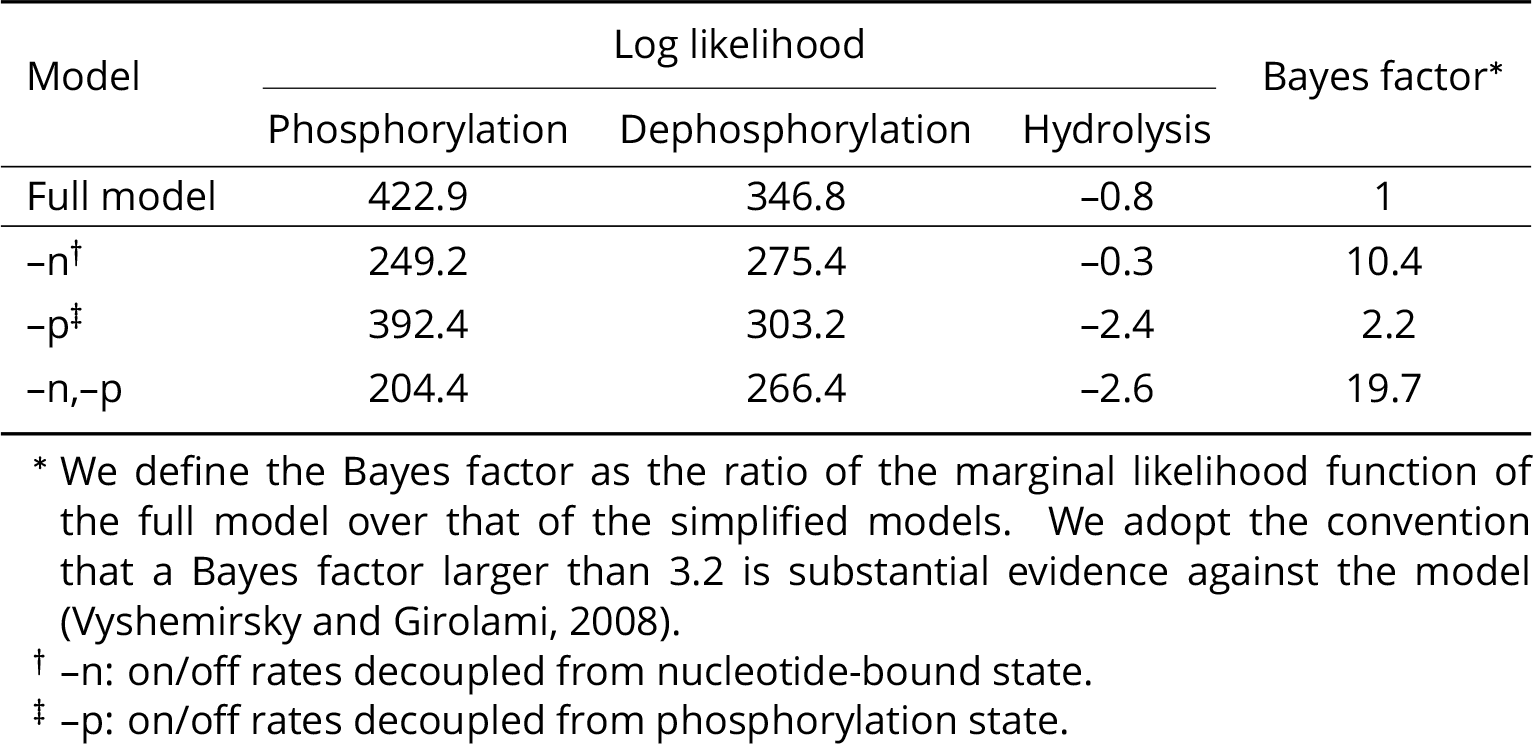
Effects of differential KaiA (un)binding kinetics

| Model | Log likelihood |  |  | Bayes factor* |
| --- | --- | --- | --- | --- |
|  | Phosphorylation | Dephosphorylation | Hydrolysis |  |
| Full model | 422.9 | 346.8 | -0.8 | 1 |
| -n<sup>†</sup> | 249.2 | 275.4 | -0.3 | 10.4 |
| -p<sup>‡</sup> | 392.4 | 303.2 | -2.4 | 2.2 |
| -n,-p | 204.4 | 266.4 | -2.6 | 19.7 |

\* We define the Bayes factor as the ratio of the marginal likelihood function of the full model over that of the simplified models. We adopt the convention that a Bayes factor larger than 3.2 is substantial evidence against the model (Vyshemirsky and Girolami, 2008).
<sup>†</sup> -n: on/off rates decoupled from nucleotide-bound state.
<sup>‡</sup> -p: on/off rates decoupled from phosphorylation state.

### Substrate competition underlies metabolic compensation

Finally, we consider the implications of the ultrasensitivity for the full oscillator. For the sake of clarity, we make a distinction in this section among three subpopulations of KaiA: the sequestered KaiA, which refers to inactive KaiA in a KaiABC complex; the active KaiA, which refers to (free or bound) KaiA not sequestered by KaiB; and the free KaiA, which is not associated with either KaiB or KaiC.

We first consider the role ultrasensitivity plays in regulating KaiA activity. It is well-established that KaiB plays an essential role in regulating KaiA during nighttime. At dusk, the buildup of KaiC D and S phosphoforms triggers the binding of KaiB to CI (Rust et al., 2007; Chang et al., 2012; Mutoh et al., 2013; Phong et al., 2013; Lin et al., 2014; Tseng et al., 2017; Snijder et al., 2017; Mukaiyama et al., 2018) and subsequently the sequestration of KaiA by CI-bound KaiB (Kageyama et al., 2006; Qin et al., 2010a). In the absence of active KaiA, the CII domain autodephosphorylates, and the KaiABC ternary complex disassembles (Snijder et al., 2017) at dawn as KaiC reaches its dephosphorylated state (Tomita et al., 2005), freeing KaiA and readying the clock for the next cycle.

This understanding of the negative feedback loop implies that the sequestration of KaiA by KaiB is a source of nonlinearity in the system that is critical for generating oscillation. Indeed, in many models of the Kai oscillator, the complete sequestration of KaiA during dephosphorylation is either a built-in or required feature for stable oscillation (e.g., Yoda et al., 2007; van Zon et al., 2007; Phong et al., 2013; Paijmans et al., 2017b). Our observation that phosphorylation is suppressed nonlinearly at low [KaiA] suggests that complete sequestration of KaiA by KaiB is not necessary to prevent phosphorylation. Indeed, there is mounting evidence that KaiB sequestration by itself is insufficient to completely inactivate KaiA during dephosphorylation. Specifically, measurements using native mass spectrometry, co-immunoprecipitation (co-IP), and native PAGE suggest that there is a significant amount of KaiA_2_C_6_ complex (Kageyama et al., 2006; Brettschneider et al., 2010) and free KaiA (Qin et al., 2010a) throughout the entire phosphorylation cycle.

To confirm that KaiA is not fully sequestered by KaiBC complexes, we used co-IP of FLAG-tagged KaiB to monitor the amount of uncomplexed KaiA in supernatant, which we interpret to be a measure of active KaiA concentration (**Figure 4**A and **Figure S7**D). The experiment shows that there is indeed a sizable amount of active KaiA in solution in the first half of the dephosphorylation stage, although the experiment does not allow us to assign absolute concentrations. Taken together, these results suggest that either the binding of KaiA to KaiBC is more labile or has lower affinity than previously assumed, or that the sequestration kinetics are slow compared to the length of the dephosphorylation stage. In either case, substantial amounts of KaiA appear to remain free of KaiABC complexes during oscillation.

**Figure 4:**
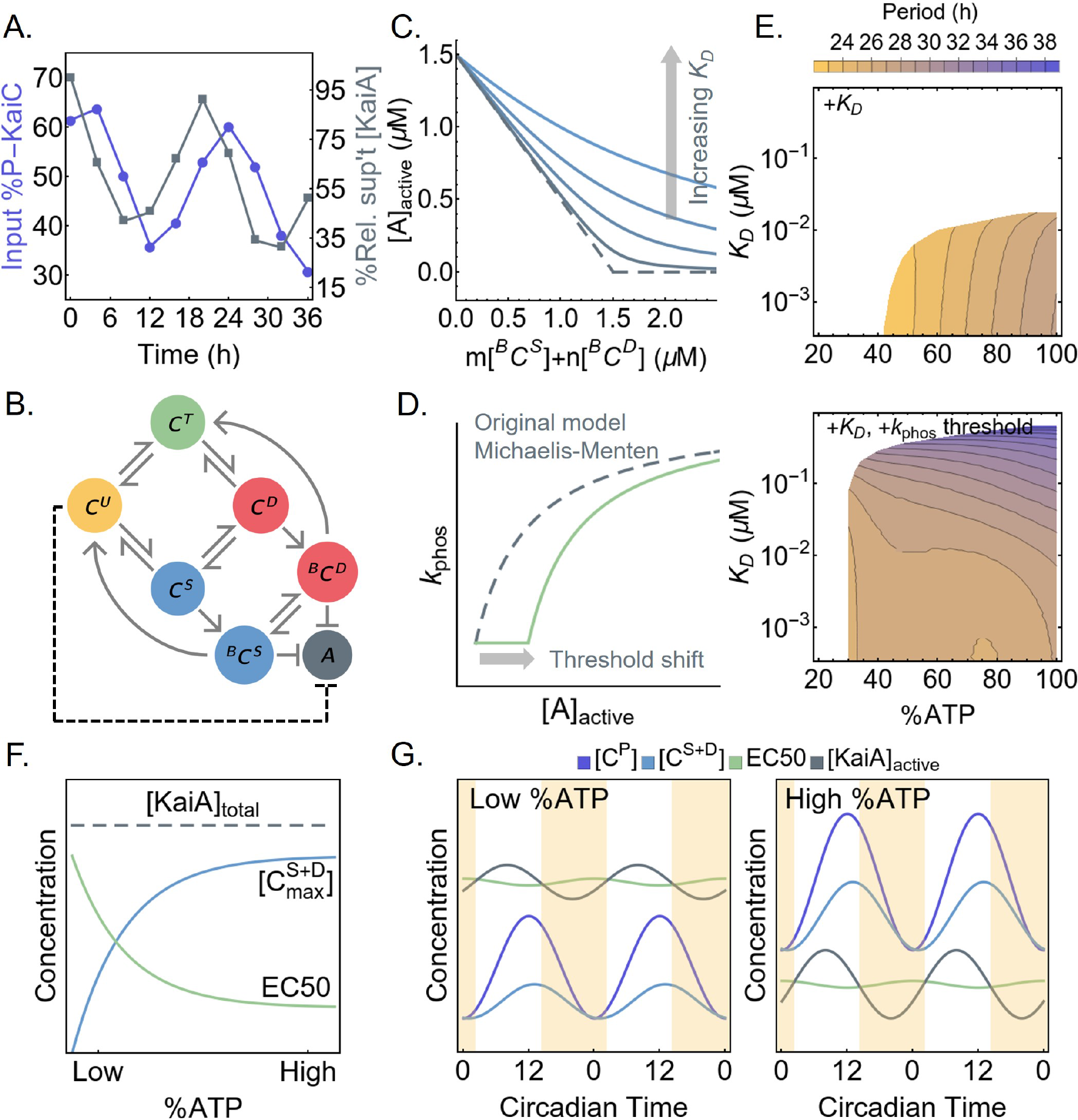
Ultrasensitivity enables metabolic compensation. A) The time series of the total input KaiC phosphorylation level (purple, left scale) and residual KaiA concentration not precipitated with KaiB-FLAG (gray, right scale). B) A schematic of the oscillator model by Phong et al. (2013). Here, ^B^C^S^ and ^B^C^D^ represent the KaiB-bound S and D phosphoforms, respectively, which can sequester KaiA. The dashed line represents the effect of introducing ultrasensitivity to the model. C) A cartoon representation of introducing a KaiA sequestration affinity, *KD*, into the Phong model. The original model has an effectively infinite sequestration affinity (dashed curve). D) A cartoon representation of introducing a KaiA threshold to the Michaelis-Menten-type phosphorylation rate constant in the Phong model. E) The period of the oscillator model as a function of %ATP and *K_D_*, a measure of KaiA sequestration affinity, without (top) or with (bottom) a phosphorylation threshold. All model simulations were done with 3.5 μM of KaiC and 1.5 μM of KaiA. White regions indicate unstable or no oscillation. F) The extent to which KaiA can be sequestered by KaiB depends on the maximal S and D phosphoform concentration, 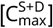, achieved over the phosphorylation cycle. The scaling of the EC50 of the phosphorylation stimulus-response function, which is a measure of the capacity of the U phosphoform to suppress KaiA activity, compensates for the scaling of 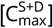 with %ATP. G) The scaling of EC50 and 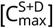 contribute to the scaling of the phosphorylation cycle size with %ATP. As %ATP increases, EC50 decreases and thus higher concentrations of the S and D phosphoforms are required to sequester active KaiA and trigger dephosphorylation. These dynamics enable increased accumulation of the T phosphoform at higher %ATP.

The fact that the phosphorylation threshold scales with %ATP suggests that ultrasensitivity may also lead to insensitivity of the period of the Kai oscillator to %ATP (Phong et al., 2013), a phenomenon termed “metabolic compensation” (Johnson and Egli, 2014). We examine this issue using a simple model of the Kai oscillator proposed by Phong et al. (2013), which we hereafter refer to as the Phong model. The Phong model explicitly keeps track of the monomer phosphorylation cycle and uses KaiB binding to the S phosphoform to generate negative feedback (**Figure 4**B). In the Phong model, the KaiA sequestration affinity is effectively infinite. In light of the co-IP experiment, we modify the model by assuming that the KaiA sequestration reaction is in a quasi-equilibrium with a dissociation constant for KaiA binding to the KaiBC complex, *K*_*D*_ (**Figure 4**C; see **SI** for mathematical details). When *K*_*D*_ is small (i.e., < 10^−3^ μM), the modified model exhibits the same robust oscillations as the original model over a large range of %ATP, but the range of %ATP that allows for stable oscillation shrinks as *K*_*D*_ increases (**Figure 4**E top), and the model is unstable when *K*_*D*_ is in the micromolar range regardless of %ATP.

In the original Phong model, the dependence of KaiC phosphorylation on KaiA is described by a MichaelisMenten-like function with no ultrasensitivity. In this scenario, a small increase in active KaiA leads to a proportional increase in phosphorylation, making the dephosphorylation phase of the clock strongly dependent on the strength of KaiB-mediated KaiA sequestration. To test if ultrasensitivity can increase the robustness of oscillations in the model, we introduce a phenomenological patch to the model in the form of an ultrasensitive KaiA threshold to the phosphorylation rate function, which varies as a function of %ATP and U phosphoform concentration (**Figure 4**D; see **SI** for mathematical details). Given that the ultrasensitivity is a result of substrate competition, this modification effectively introduces an inhibitory interaction between the U phosphoform and KaiA (**Figure 4**B, dashed arrow). This modification amounts to the assumption that the EC50 measured at steady state (**Figure 3**A) in the absence of KaiB corresponds to the active KaiA concentration required to re-enter the phosphorylation phase at the trough of the circadian oscillation. Remarkably, the resulting model can generate stable oscillations over a larger range of both %ATP and *K*_*D*_ conditions (**Figure 4**E bottom). This observation suggests that ultrasensitivity in KaiC phosphorylation plays a role in clock stability that complements the function of KaiB-dependent KaiA sequestration.

Why does ultrasensitivity in KaiC phosphorylation allow for metabolic compensation? The binding of KaiB to KaiC, and thus the sequestration and inactivation of KaiA, depends on S431 phosphorylation of KaiC (i.e., the S and D phosphoforms). At low %ATP, the maximal S and D concentrations, 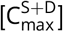, are lower (**Figure 4**F). Thus the maximal amount of KaiA sequestered by the KaiBC complex is smaller. This is problematic for the stability of the clock at low %ATP, since the active KaiA can promote premature KaiC U→T phosphorylation of some molecules, which can lead to phase decoherence, manifest as decaying oscillation (**Figure S7**E). The ultrasensitive stimulus-response that we report here implies that a finite amount of KaiA must be liberated from KaiB before there is a noticeable impact on KaiC phosphorylation. In other words, the inhibitory effect of ultrasensitivity is a synchronization mechanism. Importantly, the EC50 of the stimulus-response function scales with %ATP, such that the capacity of phosphorylation suppression by C^U^ is enhanced at low %ATP, which compensates for weaker KaiA sequestration (**Figure 4**F). This relation likely also contributes to the scaling of the phosphorylation limit cycle size with %ATP (**Figure 4**G). At higher %ATP, the EC50 is smaller and thus more KaiA needs to be sequestered to trigger dephosphorylation, which implies that higher concentrations of the S and D phosphoforms need to accumulate to enable KaiB binding. Since KaiC phosphorylation is ordered, this means that the T phosphoform concentration scales with %ATP as well.

## Discussion

In this work we undertook a data-driven kinetic modeling approach to understand the metabolic sensitivity of the KaiA-KaiC subsystem, part of the *S. elongatus* circadian oscillator. We constructed a detailed yet mechanistically agnostic kinetic model, which was fit to extensive experimental measurements of KaiC phosphorylation kinetics within a Bayesian parameter estimation framework. Approaches that are similar in spirit have been pursued in eukaryotic systems (e.g., Forger and Peskin, 2003; Locke et al., 2005; Mirsky et al., 2009; Relógio et al., 2011; Kim and Forger, 2012). However, owing to the greater complexity of eukaryotic clocks, these studies combined direct experimental measurements, cost function optimization, and hand tuning of selected parameters to account for unknown or unconstrained biochemical processes. Because the Kai system can be studied as a well-defined mixture of purified components, the participating molecular species are known, and all the parameters in the model can be treated in a consistent manner to enable objective comparison of mechanisms underlying collective oscillations.

This data-driven approach is to be contrasted with the more common hypothesis-driven, forward modeling approach, whereby a model is built to examine how features of the oscillator arise from proposed mechanisms. This hypothesis-driven approach has been employed extensively in the study of the cyanobacterial clock. These studies have revealed insights into specific aspects of the oscillator function, such as entrainment (Brettschneider et al., 2010; Leypunskiy et al., 2017), synchronization (Yoda et al., 2007; van Zon et al., 2007; Sasai, 2019), irreversibility (Cao et al., 2015), and robustness against variations in temperature (Hatakeyama and Kaneko, 2012; François et al., 2012; Kidd et al., 2015; Murayama et al., 2017), ATP/ADP concentration (Phong et al., 2013; Paijmans et al., 2017a; del Junco and Vaikuntanathan, 2019), protein copy numbers (Brettschneider et al., 2010; Lin et al., 2014; Chew et al., 2018), and environmental noise in general (Pittayakanchit et al., 2018; Monti et al., 2018). This hypothesis-driven approach is pedagogically powerful but gives little indication of the range of the parameter space consistent with a proposed mechanism, which makes it difficult to quantify the uncertainties of model predictions and validate them experimentally.

Not all parameters in our model were fully constrained by the data, as expected given the complexity of the model (Gutenkunst et al., 2007). Nevertheless, the ensemble of parameter sets still led to consistent predictions. In particular, the model revealed unexpected ultrasensitivity in KaiC phosphorylation as a function of KaiA, which we confirmed experimentally. The source of ultrasensitivity in the model is a substrate competition mechanism that arises from the differential affinity of ADP- and ATP-bound KaiC for KaiA. Previous studies have considered the importance of the differential affinity of KaiA for KaiC states but have focused on phosphorylation (Paijmans et al., 2017b; Mori et al., 2018). We note here that the ultrasensitivity in KaiC phosphorylation that we discovered acts to regulate KaiA activity at a different phase of the cycle than the ultrasensitivity in KaiB-dependent KaiA sequestration that arises from opposing S and T phosphorylations within hexamers (Lin et al., 2014). Presumably, the presence of nonlinearities and delayed feedback at multiple steps in a molecular oscillator allows the system to achieve greater robustness (Kim and Forger, 2012; Jolley et al., 2012; Dovzhenok et al., 2015; Pett et al., 2016).

We hypothesized that ultrasensitivity in KaiC phosphorylation plays a role in stabilizing the oscillator at low %ATP conditions by suppressing premature phosphorylation during the dephosphorylation stage and thus promoting phase coherence. Currently, the Kai oscillator model most robust against yet tunable by metabolic conditions appears to be that of Paijmans et al. (2017a,b). In the Paijmans model, metabolic compensation is achieved both at the hexamer and ensemble level. At the hexamer level, the onset of dephosphorylation is primarily controlled by the antagonistic effects of the T and S phosphoforms. Since fewer subunits in the T phosphoform accumulate at low %ATP, fewer subunits in the S phosphoform are needed to trigger dephosphorylation; therefore, the reduced amplitude of oscillation counteracts the slower phosphorylation rate at low %ATP. At the ensemble level, low %ATP limits the fraction of hexamers that are able to trigger dephosphorylation before the onset of KaiB-mediated delayed inhibition; this makes the dephosphorylation phase shorter, which compensates for the longer phosphorylation phase. It is worth noting that the Paijmans model is not oscillatory when %ATP reaches below 50% partly due to phase decoherence during dephosphorylation, an issue that can potentially be addressed with ultrasensitivity in KaiC phosphorylation. In our model the coupling between KaiA binding affinity and KaiC nucleotide-bound states is critical in generating ultrasensitivity, a feature that is missing in the Paijmans model. It remains an open question whether a hexameric model with no such coupling can nevertheless produce ultrasensitivity in KaiC phosphorylation (see **SI** for further comparison between this work and the Paijmans model).

In *S. elongatus*, the Kai oscillator is embedded in a transcription-translation feedback loop (Kitayama et al., 2008; Zwicker et al., 2010; Qin et al., 2010b). However, with the exception of peroxiredoxin oxidation cycles (O’Neill et al., 2011; Edgar et al., 2012), cell-autonomous circadian rhythms in eukaryotes are thought to be generated by interlocked transcription-translation feedback loops (Novák and Tyson, 2008); the cooperative autoregulation of transcription is a key source of nonlinearity and robustness in the circuit (e.g., Leloup et al., 1999; Gonze et al., 2002; Leloup and Goldbeter, 2003; Locke et al., 2005; Brown et al., 2012). Our results raise the possibility that post-translational protein modifications and protein-protein interactions may also contribute to robustness by introducing ultrasensitivity, even if these processes do not generate self-sustaining rhythms that can be decoupled from transcription. In general, it is clear that posttranslational steps such as (de)phosphorylation (Gallego and Virshup, 2007; Reischl and Kramer, 2011; Zhou et al., 2015; Fustin et al., 2018), protein degradation (Gallego and Virshup, 2007; Reischl et al., 2007), and complex formation (Kim and Forger, 2012) play an important role in eukaryotic circadian oscillators, but to our knowledge there is currently no complete experimental characterization of the stimulus-response relations of these processes.

## Materials and Methods

### Computational methods

#### Treatment of nucleotide exchange

Here we derive (4) in Results. The nucleotide exchange process is in principle a two-step reaction that includes an *apo* intermediate state of KaiC, i.e.,

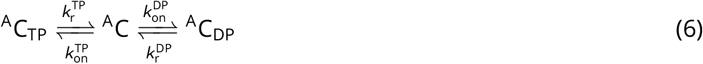

where we have omitted the free ATP and ADP from the chemical equation. Here, 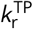 and 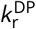 are the dissociation rate constants and 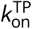 and 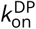 are the binding rate constants for ATP and ADP, respectively. Since KaiC requires nucleotides for hexamerization (Hayashi et al., 2003, 2006; Mutoh et al., 2013), the *apo* state of KaiC is presumably both thermodynamically and kinetically unstable in the presence of saturating amount of nucleotide (5 mM in our experiments). Therefore, under the assumption that the KaiC *apo* state is in a quasi-steady state throughout the reactions, we can eliminate the *apo* state and model nucleotide exchange as a one-step reaction

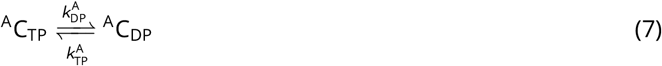

where

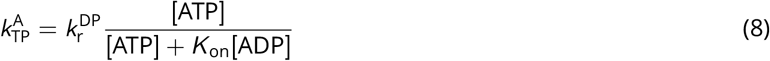

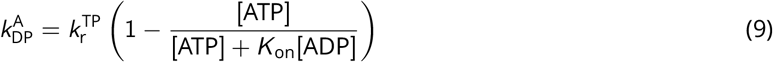

and 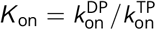 is a ratio of the two nucleotide binding rate constants.

We make two further simplifying assumptions. First, we assume the on rates are completely diffusion controlled and are thus the same for ATP and ADP, which allows us to set *K*_on_ = 1. Second, based on fit results (**Figure S4**F) showing that the posterior for 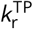 has a long tail to negative infinity in log space, we follow the approach proposed by Transtrum and Qiu (2014) and set 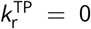; i.e., the dwell time of ATP-bound states are sufficiently long such that a bound ATP cannot be released without first giving up its γ-phosphate group. This assumption implies that the only ways for KaiC to enter an ADP-bound state are through hydrolysis and phosphorylation, and solution ADP has no effect on the system except to slow down the ADP to ATP exchange process. With these two assumptions, we eliminate (9) and (8) reduces to (4).

#### Model parameterization

In Results, we introduced a model parameterization scheme in which rate constants for phosphotransfer, nucleotide exchange, KaiA (un)binding, and ATP hydrolysis reactions depend on the participating molecular species. Although we use this independent-rate scheme to interpret the model, including computing the sensitivity ODEs, during the fitting itself we represent species-dependent effects by modifying each of the six basic rate constants (*k*_p_, *k*_d_, *k*_a_, *k*_b_, 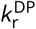, and *k*_h_) by multiplicative Δ*k* factors. For example, the KaiA dissociation rate 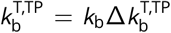 is represented by the product between a base rate *k*_b_ and a modifier 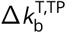 (compare **Figure S1** and **Figure S10**). The multiplicative-factor scheme introduces 38 Δ*k* parameters. Because of the requirement for detailed balance (see below), only 34 of these parameters are free; these free parameters are listed on **Table 2**. The advantage of the multiplicative parameterization scheme is that it facilitates *ℓ*^1^ regularization, discussed below.

**Table 2:**
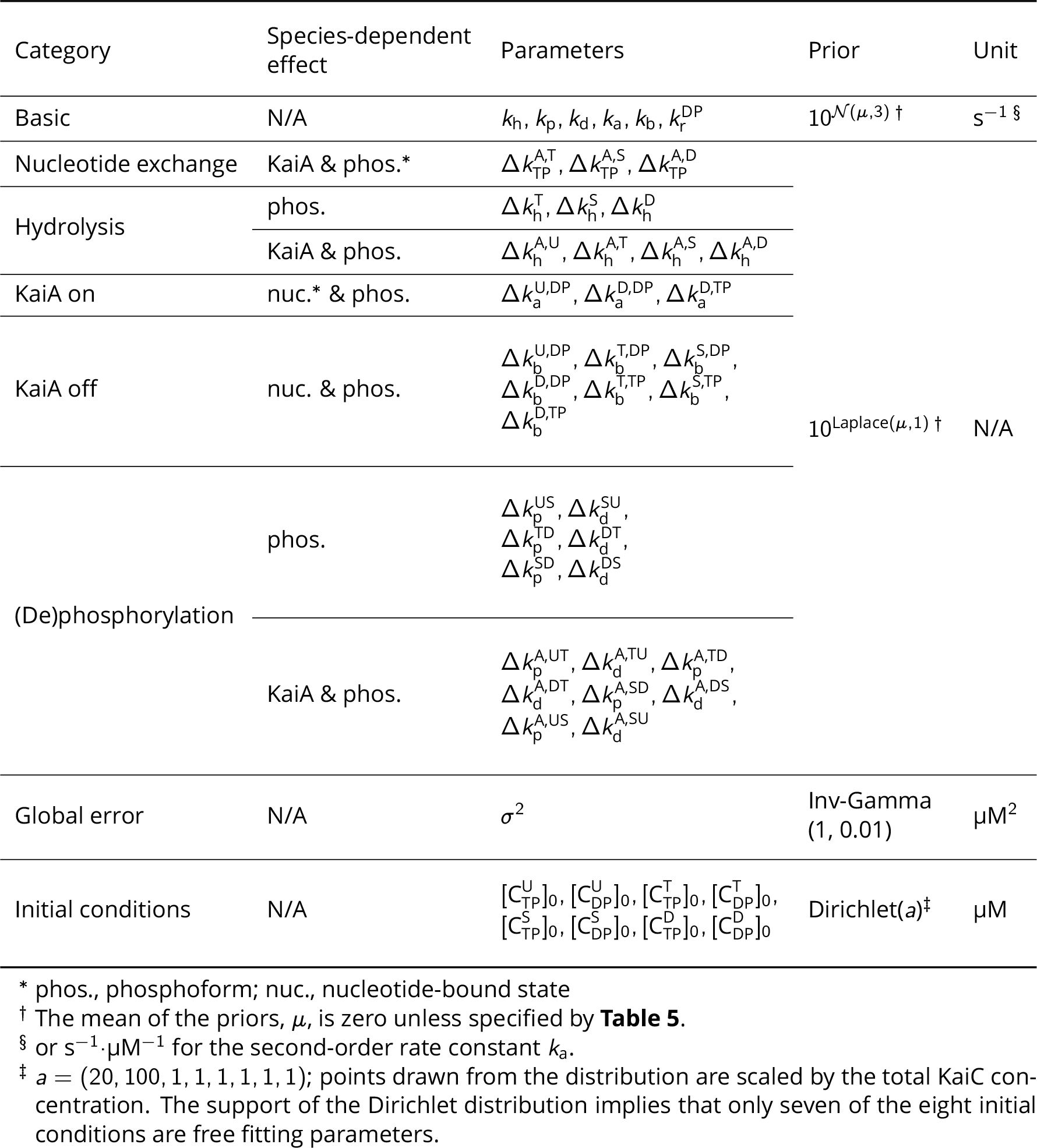
Full model parameters and their priors

#### Detailed balance

All elementary reactions, except ATP hydrolysis and nucleotide exchange, are assumed to occur in equilibrium, and thus the free energy change over each reversible cycle must be equal to zero.

In practice, this means that the product of all rate constants in the forward direction of each cycle listed on **Table 3** must be equal to that in the reverse direction (see also **Figure S10**A). This introduces an additional algebraic constraint for each such cycle, which is used to eliminate one free Δ*k* parameter. In total, one can eliminate four such arbitrarily chosen parameters.

**Table 3:**
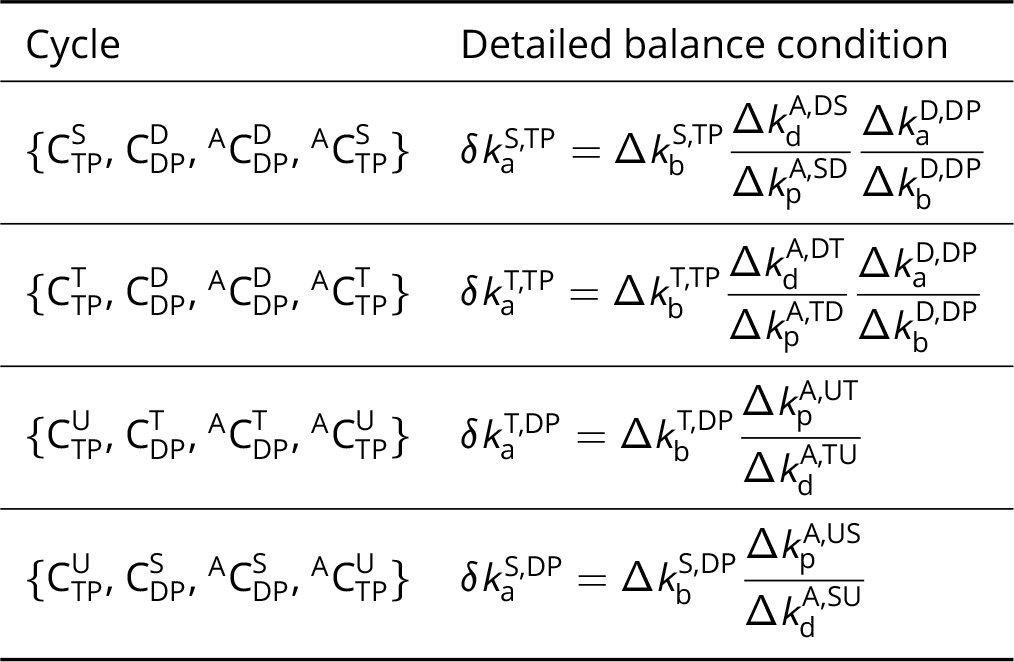
Detailed balance conditions

#### Fitting data set

To constrain the model parameters, we collected experimental measurements that characterized different aspects of the KaiA-KaiC subsystem, which are summarized in **Table 4**.

**Table 4:** Fitting data set

| Measurement (source) | Temperature (°C) | [KaiA]* (μM) | %ATP | Time points | Phosphoform |
| --- | --- | --- | --- | --- | --- |
| Phosphorylation (this work) | 30 | 0.375, 0.75, 1.50, 3.00, 4.50, 6.00 | 10, 25, 100 | 8 | U, T, D<sup>†</sup> |
| Dephosphorylation (Rust et al., 2011) | 30 | 1.4 | 100 | 21 |  |
| ADP production (Terauchi et al., 2007) | 30 | 1.2 | 100 | 1 | N/A |
| KaiA on/off rates (Kageyama et al., 2006) | 25 | Variable | 100 | N/A | Likely U |
| KaiA dwell time (Mori et al., 2018) | 25–28 | 1.0 | N/A | N/A | T, S, D<sup>‡</sup> |

\* We report here on the KaiA monomer concentration. However, since KaiA functions as a dimer, all KaiA concentration is divided by two in the models.
<sup>†</sup> The conservation of mass constraint implies that one of the four phosphoforms is not a free state variable. We have chosen the S phosphoform to be the constrained state variable.
<sup>‡</sup> Phosphomimetic mutants

The dephosphorylation reaction taken from Rust et al. (2011) constrains the dephosphorylation rates and the ATP hydrolysis rates of KaiC in the absence of KaiA, because the model structure dictates that dephosphorylation requires alternating phosphotransfer and hydrolysis reactions. There is currently no direct measurement of CII hydrolysis rate in the presence of KaiA. However, the maximum ADP production rate of KaiC in the presence of 1.2 μM of KaiA was reported to be 29.8 ± 5.1 KaiC^−1^ day^−1^ (Terauchi et al., 2007), which we take as an upper bound on the average CII hydrolysis rate in phosphorylation reactions for all [KaiA] = 0.375, 0.75, and 1.50 μM conditions.

Because the phosphorylation reactions were measured in the presence of varying %ATP and higher [ADP] inhibits phosphorylation by slowing down nucleotide exchange (see equation 4), they provide indirect constraints on the nucleotide exchange rate. Similarly, because the reactions were measured in the presence of varying [KaiA] conditions, they provide direct constraint on the phosphorylation rates of KaiC with and without KaiA, as well as the KaiA binding affinity, i.e., the ratio of KaiA on/off rates. Although there are direct experimental measures of KaiA binding and dissociation (Kageyama et al., 2006; Mori et al., 2018), these results cannot be directly mapped onto model rate constants. This is primarily because the KaiC nucleotide-bound fractions are not reported in these experiments, or, in the case of phosphomimetic mutants, it is unclear if the mutations affect nucleotide binding affinities. As a consequence, the experimental constraints on KaiA on/off rates enter through the priors rather than the likelihood function, in contrast to the other data (see below).

#### Initial conditions

For each phosphorylation reaction in **Table 4**, we solve the ODE model with the corresponding [KaiA] and %ATP condition. The predicted phosphoform composition, as well as the ATP hydrolysis rate when appropriate, is compared to the experimental measurements in the Bayesian parameter estimation formalism described below. However, since the experimental data do not resolve the initial conditions for all 16 KaiC species in the model, we have chosen to directly estimate the initial concentrations as free model parameters. We take *t* = 0 to be the time point at which KaiA is mixed with KaiC, and thus all eight KaiA-bound KaiC states are assumed to have zero concentration at the onset of the experiment. Because total KaiC concentration is conserved, this introduces seven additional free parameters (see **Table 2**).

For the dephosphorylation reaction, we do not estimate the initial conditions. To mimic the way the experiment was done, the dephosphorylation reaction is simulated in two stages. In the first stage, we assume that 3.4 μM of dephosphorylated KaiC is phosphorylated in the presence of 1.3 μM KaiA and 100% ATP for 20 h. Since the protein is initially dephosphorylated, we assume that 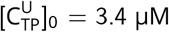 while the concentrations of all other KaiC species are set to zero. In the second stage, we simulate the autodephosphorylation reaction after KaiA pull-down, which corresponds to eliminating all free KaiA as well as KaiA-bound KaiC species from the simulation. The amount of KaiC lost in the pull-down experiment was not reported in the original experiment (Rust et al., 2011). We therefore make the assumption that the amount of KaiC lost in the pull-down experiment in the simulation is exactly equal to that in the experiment for the purpose of computing the likelihood function.

#### Bayesian parameter estimation

We directly fit numerically integrated ODEs to experimental data in the Bayesian parameter estimation framework (Wasserman, 2000). The best fit model parameters, 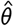, are obtained from the maximum a posteriori estimator:

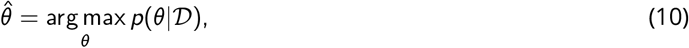

where 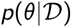 is the posterior distribution of the parameters *θ*, conditioned on the training data set 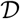. Using Bayes’ theorem, the posterior distribution can be written as

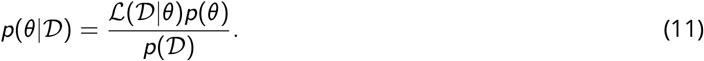

Here, *p*(*θ*) is the prior distribution, which represents subjective belief in the model parameters *θ* prior to experimental input; 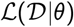 is the likelihood function, which represents a probabilistic model of the experimental data set 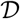 given a particular model 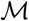 (implicit in the formulas) that depends on the parameters *θ*; 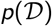 is the evidence, which is analogous to the partition function in statistical mechanics. Note that the evidence 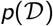 does not depend on the parameter choice, and is thus an irrelevant constant for the purpose of parameter estimation. The specific choices for the functional forms of the likelihood function and priors are discussed further below.

#### Model priors

The priors for all model parameters used in Bayesian parameter estimation are given in **Table 2**. Here, the choice of the prior distributions is primarily motivated by the need for regularization (see below). In addition, as discussed above, the experimental measurements on KaiA binding kinetics are incorporated into the priors rather than the likelihood function (**Table 5**). Note that all the rate constants and their multiplicative factors are estimated in the log space (base 10). This ensures that all rate constants are positive.

**Table 5:** Priors incorporating KaiA on/off constraints

| Parameter | Prior mean ($\mu$) | Experimental measurements | Source |
| --- | --- | --- | --- |
| $k_a$ | $\log k_{a,\text{exp}}$ | $k_{a,\text{exp}} = 0.0279 \text{ s}^{-1} \cdot \mu\text{M}^{-1}$ | Kageyama et al. (2006) |
| $k_b$ | $\log k_{b,\text{exp}}$ | $k_{b,\text{exp}} = 0.0663 \text{ s}^{-1}$ | |
| $\Delta k_b^{\text{T,DP}}, \Delta k_b^{\text{T,TP}}$ | $-\log \tau_{b,\text{exp}}^T k_{b,\text{exp}}$ | $\tau_{b,\text{exp}}^T = 1.0 \text{ s}$ | Mori et al. (2018) |
| $\Delta k_b^{\text{S,DP}}, \Delta k_b^{\text{S,TP}}$ | $-\log \tau_{b,\text{exp}}^S k_{b,\text{exp}}$ | $\tau_{b,\text{exp}}^S = 0.43 \text{ s}$ | |
| $\Delta k_b^{\text{D,DP}}, \Delta k_b^{\text{D,TP}}$ | $-\log \tau_{b,\text{exp}}^D k_{b,\text{exp}}$ | $\tau_{b,\text{exp}}^D = 0.26 \text{ s}$ | |

#### *ℓ*^1^ regularization

As model complexity grows, the constraint of experimental data on the underlying mechanism weakens. To address this problem, we impose sparsity on the species-dependent effects (i.e., the Δ*k* factors) using *ℓ*^1^ regularization (Tibshirani, 1996). This is accomplished in the Bayesian parameter estimation framework by using a Laplace prior centered at zero in the log parameter space (or one in the real space). Intuitively, the Laplace prior imposes sparsity by forcing the (marginalized) posterior distribution for each log Δ*k* to peak at zero unless there is experimental evidence in the fitting data set to suggest otherwise. Since the Δ*k*s are multiplicative factors modifying the six basic rate constants, log Δ*k* = 0 implies that the species-dependent rate is identical to that of the base rate. This method is directly analogous to the use of the lasso estimator in the context of linear least squares models. To see this, consider the Laplace distribution

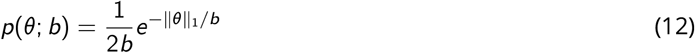

where ‖*θ*‖_1_ is the *ℓ*^1^-norm of *θ*. Then from (11) the negative log-posterior distribution becomes

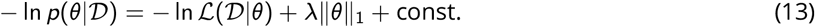

where λ = 1/*b*. In a linear model *Y* = *Xβ* + *ϵ* where *Y* is the response vector, *X* is the design matrix, *β* is the parameter vector, and *ϵ* is the error vector, the negative log-likelihood function reduces to the sum of squares 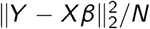, where *N* is the number of dependent variables. Thus, maximizing the posterior is equivalent to minimizing the sum of squares with an *ℓ*^1^ penalty, which is the lasso estimator.

#### Likelihood function

To determine the functional form of the likelihood function, consider a kinetic experiment where measurements on some observables *y* are made at a set of time points {*t*_*i*_} with uncertainties {*σ*_*i*_}. If we assume that the experimental errors *σ*_*i*_ are independent and normally distributed, then the likelihood function is given by

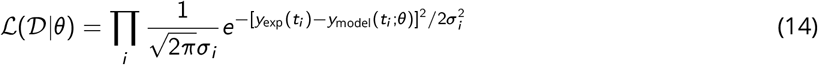

In other words, the likelihood function gives the probability for observing a given data set provided that the model prediction is true. In practice, all posterior evaluations are done in the log space (base *e*) for numerical stability. Thus, taken together, (11) can be rewritten as,

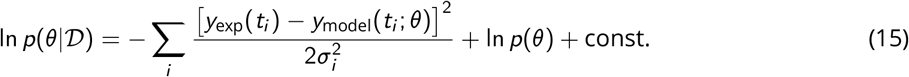

For the sake of simplicity, we assume that there is a single global error, *σ*, for all (de)phosphorylation measurements, which is then estimated during fitting as a hyperparameter (see **Table 2**).

The choice of the Gaussian likelihood function applies to all (de)phosphorylation data sets, but not the hydrolysis constraint, which only provides an upper bound on the average hydrolysis rate per day (Terauchi et al., 2007). Therefore, for the hydrolysis data a “half harmonic” is used as the log-likelihood function:

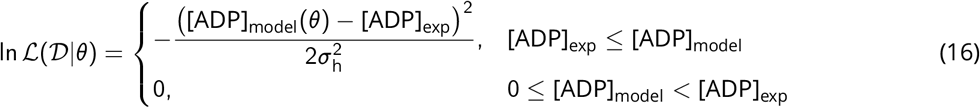

The total amount of ADP produced by the model during a phosphorylation reaction over 12 h, [ADP]_model_, is given by the sum of all P_*i*_ production over time plus all ADP-bound KaiC species at *t* = 12 h.

The log-likelihood values from the appropriate phosphorylation, dephosphorylation, and hydrolysis reactions are added together to determine the log-likelihood of the data set for each given model parameter choice. Since the phosphorylation data set is much larger than the dephosphorylation data set, the fitting procedure tends to favor fitting the phosphorylation data set at the expense of fitting the dephosphorylation data set. To overcome this problem, the log-likelihood function for the dephosphorylation reaction is multiplied by a factor of 4 to increase the weight of the dephosphorylation data points.

#### Model fitting procedure

To determine the posterior mode and the uncertainties associated with the estimate, we employ a heuristic combination of ensemble MCMC sampling and numerical optimization methods (**Figure S3**A). This fitting procedure can be divided into four steps that are analogous to those in a genetic algorithm:

1. Initialization. An ensemble MCMC method evolves a set of random walkers (i.e., parameter sets) simultaneously; we thus begin by drawing 224 walkers from the prior distribution *p*(*θ*) and use these walkers for simulated annealing (Kirkpatrick et al., 1983; Kirkpatrick, 1984). In annealing, instead of sampling 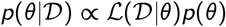 (in the log space), a flattened distribution 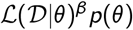 is sampled with an annealing schedule of *β* = 0.3, 0.4, 0.5, 0.6, 0.7, 0.8, 0.9, 1.0. Note that instead of letting *β* → ∞, the simulation ends with the target distribution at *β* = 1.0. Each temperature is sampled over 20,000 steps.
2. Selection. The fitnesses of the walkers are determined by their log-posterior values (equation 15). 10 walkers from the best 300 parameter sets sampled in the previous step are chosen and subjected to numerical optimization to find the nearby local maximums, which are then used to seed a sampling run in the next step. In the spirit of elitist selection, the best walker is always included for the next generation.
3. Recombination and mutation. The initial walkers for the sampling run are generated by adding a Gaussian noise 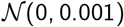 to the 10 optimized walkers, and the number of initial walkers centered around each optimized walker *θ*_*j*_ is given by

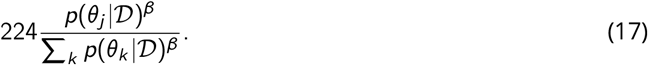 That is, the proportion of the initial walkers generated from each optimized walker is weighted by its posterior value with a temperature factor of *β* = 0.6; the temperature factor is chosen to allocate more walkers to optimized walkers with lower posterior values. The sampling run consists of 50,000 steps. Note that the purpose of the Gaussian noise is to ensure that the proposal distribution is valid for any pair of walkers for the ensemble MCMC method (see below), rather than to control the mutation strength, as is done in evolution strategy (Beyer and Schwefel, 2002).
4. Termination. The best walkers from the sampling run are compared to the optimized walkers. If the best walkers do not escape to new local maximum(s) with higher posterior value(s), then the procedure is terminated after an additional 50,000 sampling steps. If, however, new local maximum(s) are discovered during sampling, the algorithm loops back to the selection step. This process is repeated until no better local maximum is discovered at the end of sampling. Unless otherwise specified, only the last 30,000 sampling steps (downsampled every 100 steps) are used for post-analysis.

In general, the number of walkers in ensemble MCMC needs to be larger than the number of free parameters; here the number 224 is chosen to optimize parallel performance on a local computer cluster (8 nodes × 28 CPU cores/node).

We found that this procedure outperformed either ensemble MCMC or numerical optimization by itself (compare **Figure S3**A and B). For the full model we ran this procedure three times to assess the reproducibility of the fit (see **SI** for further discussion).

#### Markov chain Monte Carlo

One major challenge in efficient MCMC sampling in systems biology is that the target distributions are often poorly scaled. In the context of ODE kinetic modeling, this means that different reaction rates and their associated uncertainties can be separated by several orders of magnitude. This is almost certainly true for the KaiA-KaiC subsystem because, among other things, the experimentally measured rates of KaiA binding and dissociation are much faster than the ATP hydrolysis rate of KaiC. Without *a priori* knowledge of the natural time scales, conventional MCMC schemes are inefficient in such sampling problems, because only very small displacements are accepted at appreciable rates. In this work we employ an ensemble MCMC method developed by Goodman and Weare (2010). The advantage of the Goodman-Weare algorithm is that it is affine invariant, which means that it performs equally well for isotropic and poorly scaled measures, providing that the two can be related by a linear transformation of the coordinate system. This appears to be the case for the present problem since the Goodman-Weare algorithm vastly outperforms a standard Metropolis-Hastings scheme with a (preconditioned) Gaussian proposal distribution (**Figure S3**B).

In brief, the Goodman-Weare algorithm evolves an ensemble of walkers, rather than a single one. At each step, individual walker positions are updated sequentially. For a given walker *θ*_*k*_ at step *τ*, a walker *θ*_*j*_ is drawn randomly from the rest of the ensemble and a new position, *η*, on the line connecting *θ*_*k*_ and *θ*_*j*_ is proposed by a “stretch move”

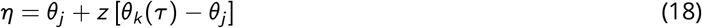

where *z* is a random number drawn from the distribution

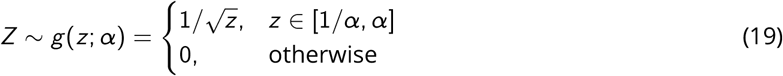

The “stretch factor” *α* is a tunable parameter that controls the step size. In an *N*-dimensional parameter space, the new walker *η* is accepted with the probability

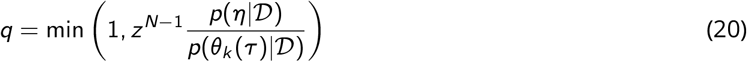

which guarantees that the scheme obeys detailed balance and thus converges to the target distribution 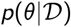 as *τ* → ∞. Note that no derivative of the posterior distribution is required to draw from the proposal distribution. In this work we use *α* = 1.1, which gives an average acceptance rate of 47% in steps 3 and 4 of the fitting procedure.

#### Numerical optimization

The numerical optimization method used in this work is a modified version of Powell’s method (Powell, 1964; Press et al., 2007). Briefly, given an initial guess and direction set, which is usually the Cartesian coordinate set, Powell’s method performs a line search to minimize the objective function, here 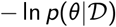, sequentially along each vector in the direction set. The direction set is then updated by replacing the direction of largest decrease in the objective function in the current iteration with the displacement vector from the estimated minimum at the beginning to that at the end of the line minimizations, provided that certain technical conditions are met to avoid the build-up of linear dependence in the direction set. This process is repeated until a convergence threshold is met. Note that unlike the original method, the modified Powell’s method does not guarantee that the vectors in the direction set are mutually conjugate.

Similar to the Goodman-Weare algorithm, Powell’s method is derivative-free. For the current system Powell’s method converges faster than the Nelder-Mead method (Nelder and Mead, 1965), another commonly used derivative-free method, although the Nelder-Mead method appears less prone to becoming trapped in local metastable states (**Figure S3**C).

#### Software implementation

The fitting procedure is implemented in an in-house Python (Oliphant, 2015) script that interfaces with several existing Python modules: numerical integration of the model ODEs is done using the odespy package (Langtangen and Wang, 2015) with the BDF method; the Goodman-Weare algorithm is implemented in emcee (version 2.2.1) (Foreman-Mackey et al., 2013); Powell’s method and the Nelder-Mead method are implemented in scipy (version 1.2.1) (Eric Jones et al., 2001). The derivative evaluation step in ODE integration is accelerated using numba (Lam et al., 2015), and the script is parallelized using mpi4py 2.0.0 (Dalcín et al., 2005, 2008; Dalcin et al., 2011).

The most computationally expensive step in the fitting procedure is the MCMC sampling, because each move requires multiple ODE evaluations to compute the posterior function. With 224 walkers and 8 nodes (each with 28 Intel E5-2680v4 2.4GHz cores), the speed of MCMC sampling is 46,000 steps/hour.

#### Model comparison

In the preceding sections all definitions of probability distributions implicitly assume that there is a model 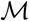 with a well-defined functional form, whose parameters *θ* need to be determined. For the sake of model comparison, we make this assumption explicit and rewrite (11) as

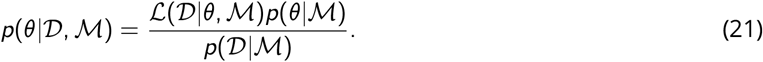

To compare two models 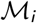 and 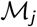, we need to compare the posterior probabilities for each model, usually in the form of their ratios

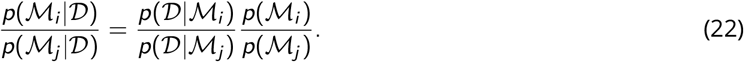

Assuming that we have no prior preference for any model, the ratio becomes the Bayes factor

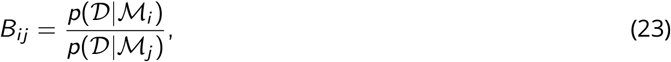

which we adopt as the metric for model comparison.

The primary difficulty in computing the Bayes factor is thus estimating the evidence, or the marginal likelihood function, for each 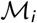. There are several methods for computing the evidence (Vyshemirsky and Girolami, 2008). Here we derive a formula compatible with the ensemble MCMC scheme that is directly analogous to free energy perturbation (Zwanzig, 1954). For the sake of simplicity, we drop the model index *i* from this point on. First, note that for a given model 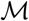,

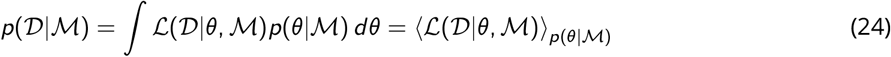

where the first equality follows from the law of total probability and the second equality assumes that the prior 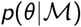 is normalized (as a probability density function of *θ*). Equation (24) suggests that the marginal likelihood function can be computed by estimating the average of the likelihood function 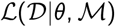 against the prior. Using MCMC to estimate this integral is inefficient since there is very little overlap between the likelihood function and the prior for the models of interest. Instead, we define

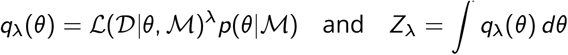

for 0 ≤ λ ≤ 1 and then note that (24) can be recast as

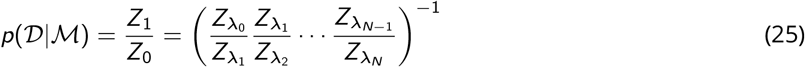

for 0 = λ_0_ < λ_1_ < … < λ_*N*_ = 1, and the λs are chosen to allow for sufficient overlap between successive *q*_λ_(*θ*)s. Each fraction in (25) is given by

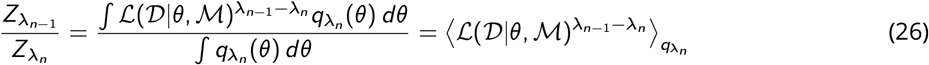

Therefore,

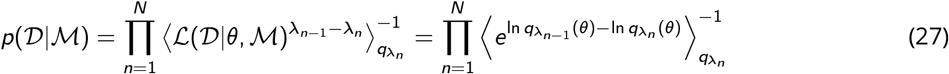

where the averages 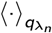 can be approximated with MCMC. Equation (27) is a version of the free energy perturbation formula. Note that (27) requires that the likelihood function 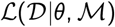 be properly normalized (as a probability density function of 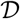), but does not require the prior 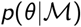 to be normalized, as any missing normalization constant cancels out in each term of the product.

For each simplified model in **Table 1** and **Table S1**, the ensemble of walkers from the last time step of the model fitting procedure is used to initialize an MCMC sampling run with λ = 1.00. The lambda value is reduced by 0.01 at each subsequent stage until λ reaches 0.01. Each stage is sampled for 2,000 time steps using the Goodman-Weare ensemble sampler Goodman and Weare (2010). Only the last 1,000 time steps from each stage is used to compute the ensemble average in (27). The Bayes factors are then computed as the ratios of the evidence for the full model to each simplified model.

#### Refitting

The steady-state KaiC phosphorylation measurements (**Figure 3**B and **Figure S8**B) are fit to the full model using Powell’s method, starting from the best fit based on the training data set. The priors on the kinetic parameters (**Table 2**) are centered on the best fit values, so that the refit model can be interpreted as the “minimal” perturbation to the best fit that enables agreement with the steady-state measurement.

#### Curve fitting

The experimentally determined stimulus-response relations for KaiC S431A (**Figure 3**C and F) are fit to the simple inhibitor ultrasensitivity scheme described in Box 5 of Ferrell and Ha (2014b). Using their notation, the amount of phosphorylated protein substrate (%*XP*) as a function of kinase concentration ([*K*]) is given by

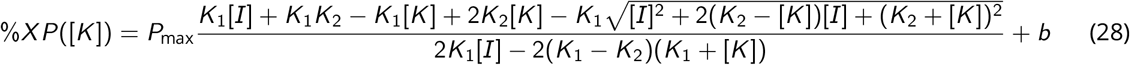

Here, *P*_max_, [*I*], *K*_1_, *K*_2_, and *b* are free model parameters. Unlike the Hill function, EC50 is not an explicit parameter of (28) and thus needs to be determined numerically. Note that (28) can be reduced to a right-shifted hyperbolic function as *K*_2_ → 0:

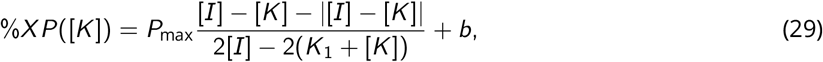

which is equivalent to a threshold-hyperbolic stimulus-response function,

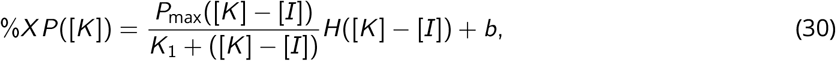

where *H* is the unit step function.

Stimulus-response relations are fit using the NonlinearModelFit function in Mathematica 12.0. We stress here that the curve fits are purely phenomenological and are thus not intended to be interpreted in terms of the biochemical assumptions underlying the model.

### Experimental methods

#### Protein expression and purification

KaiA, KaiB, KaiB-FLAG, and KaiC were expressed and purified as previously described (Phong et al., 2013) with two modifications to the protocol: anion exchange chromatography was performed using HiTrap Q columns (GE Healthcare), and KaiC was purified using Ni-NTA affinity chromatography followed by size-exclusion chromotography, omitting the anion exchange step. The expression, purification, and 6-iodoacetofluorescein (6-IAF) labeling of KaiB K25C mutant as a fluorescence reporter in the plate reader assay is described in Leypunskiy et al. (2017). All mutants of KaiC were constructed using QuikChange II XL Site-Directed Mutagenesis Kit (Aligent). For the KaiC AA and EE mutants, the His-tags were not cleaved during purification; this ensures that these mutant proteins have shifted mobility in SDS-PAGE, allowing their bands to separate from those of KaiC S431A.

The U-[^15^N] labeled N-terminal (residues 1–135) and C-terminal (residues 181–284) domains of KaiA were expressed were expressed in BL21(DE3) *E. coli* (Novagen) in minimal (M9) media supplemented with ^15^N-enriched NH_4_Cl. For expression of C-terminal domain, M9 media enriched with ^15^N-NH_4_Cl was prepared using 98% deuterated water (D_2_O). The proteins were purified by Ni-NTA affinity chromatography followed by size-exclusion chromatography using Superdex 75 1660 prep grade column, as described previously (Chang et al., 2011, 2012; Tseng et al., 2014). N-terminal KaiA eluted as ~15 kDa monomer (Vakonakis et al., 2001), while C-terminal KaiA eluted as ~23 kDa homodimer (Vakonakis and LiWang, 2004).

GFP was expressed as N-terminal 6xHis-tag fusion from the pET28a plasmid in the BL21 (DE3) strain of *E. coli*. Harvested cells were sonicated for lysis and clarified lysate was loaded onto a HisTrap FF column (GE Healthcare). The His tag was cleaved by overnight incubation at 4 °C with SUMO protease (Invitrogen), after which the sample was loaded again onto a HisTrap FF column to recover the cleaved products. The cleaved proteins were further purified on a 5 mL HiTrap Q HP column (GE Healthcare) and then a Superdex 200 10/300 GL size-exclusion column. The eluted fractions were concentrated in a sample buffer (20 mM HEPES [pH 7.4], 150 mM KCl, 2.5 mM MgCl_2_, 2 mM DTT), aliquoted, and snap frozen in liquid nitrogen for storage in −80 dC.

#### In vitro clock reactions

All in vitro clock reactions were done in the standard reaction buffer (20 mM Tris-HCl [pH 8], 150 mM NaCl, 5 mM MgCl_2_, 0.5 mM EDTA, 10% glycerol, 50 μg/mL Kanamycin). KaiC concentration was 3.5 μM in all experiments unless otherwise specified; KaiB concentration was 3.5 μM for all oscillatory reactions, and 6-IAF-labeled KaiB K25C concentration was 0.2 μM for plate reader assays. KaiA concentration and %ATP were determined by each individual experiment, while the total nucleotide concentration (i.e., [ATP] + [ADP]) was held constant at 5 μM. Phosphorylation kinetics was resolved using SDS-PAGE on 10% acrylamide gels (37.5:1 acrylamide:bis-acrylamide) run for 4.5 h at 30 mA constant current at 12 °C; the gels were stained in SimplyBlue SafeStain (Invitrogen) and then imaged using Bio-Rad ChemiDoc Imager. The oscillatory reactions (**Figure 2**D) were also monitored using the plate reader assay described in Leypunskiy et al. (2017).

#### NMR spectroscopy

A Bruker 600 MHz AVANCE III spectrometer equipped with a TCI cryoprobe was used for all of the NMR experiments of the N- and C-terminal domains of KaiA (**Figure S2**). Chemical shifts were referenced to internal 2,2-dimethyl-2-silapentane-5-sulfonate (10 μM). Data were processed using NMRPipe and visualized using NMRDraw (Delaglio et al., 1995). NMR samples were prepared with 100 μM monomer concentration of protein in 20 mM Tris-HCl [pH 8], 150 mM NaCl, 5 mM MgCl_2_, and 5% D_2_O buffer. All experiments were performed at 30 °C. Samples were incubated with 1mM ATP or ADP, when needed, for 30 minutes before spectral measurement.

#### Immunoprecipitation

Immunoprecipitation of KaiB-FLAG and associated protein complexes in a clock reaction (**Figure 4**A) was done as previously described (Phong et al., 2013). The supernatant was analyzed by SDS-PAGE on 4–20% Criterion TGX Stain-Free Precast Gels (BioRad) and stained with SYPRO Ruby (BioRad). 1.5 μM GFP was added to the reaction mixture at the beginning of the time course and serve as an internal standard to correct for changes in protein concentration due to handling. The relative supernatant KaiA concentration was determined as a ratio of KaiA band intensity in each lane to the GFP band intensity and is normalized as percentage of the largest ratio in the time course (**Figure S7**D).

## Acknowledgments

We thank Connie Phong and Haneul Yoo for their protein samples, and Jonathan Weare for helpful discussions. This work was supported by National Institutes of Health awards GM109455, GM107369, GM107521, and EY025957, Department of Energy Office of Advanced Scientific Computing Research contract DE-AC02-06CH11347 and award DE-SC0014205, and a Howard Hughes Medical Institute Faculty Scholarship (to MJR). AL was also supported by the Center for Cellular and Biomolecular Machines at University of California, Merced (NSF Grant HRD-1547848). Computations were performed on resources provided by the University of Chicago Research Computing Center, and the Extreme Science and Engineering Discovery Environment (Towns et al., 2014) (NSF Grant ACI-1548562) Bridges (PSC) computing nodes through allocation TG-MCB180007.

## Supplementary Information

### Additional biochemistry of the KaiC (de)phosphorylation reactions

In this work we construct a general model of the KaiA-KaiC subsystem based on a set of assumptions of basic clock biochemistry; that is, KaiC is an ATPase and a reversible phosphotransferase with two phosphorylation sites at S431 and T432, while KaiA is a nucleotide-exchange factor that promotes the exchange of bound ADP for ATP in CII nucleotide-binding pockets. Through model fitting, we demonstrate that this set of assumptions is sufficient to explain the (de)phosphorylation kinetics of KaiC and its dependence on %ATP and [KaiA]. Our results, however, do not imply that the model is biochemically exhaustive; in this section and the next we briefly discuss aspects of KaiC enzymology that we do not consider in the model.

First, the current model does not account for the CI domain. The CI domain of KaiC is required for the hexamerization of *S. elongatus* KaiC (Hayashi et al., 2004b, 2006) and its ATPase activity is necessary for the allosteric transition into the dephosphorylation phase of the circadian cycle (Phong et al., 2013; Tseng et al., 2017). However, the hydrolysis state of the CI domain has no significant effect on the CII (de)phosphorylation reactions (Phong et al., 2013), and in the current study we are not concerned with KaiB-dependent processes. Therefore the current model does not keep track of the CI hydrolysis state or the allosteric coupling between the CI and CII domains.

Second, the model does not explicitly consider the function of Mg^2+^. The presence of Mg^2+^ is required for the assembly of the KaiC hexamer (Hayashi et al., 2006; Mutoh et al., 2013), and computational analyses indicate that release of Mg^2+^ independent of the bound nucleotide is highly energetically unfavorable (Hong et al., 2018). Given these results, we assume that Mg^2+^ and nucleotide cannot act independently of each other, and the model implicitly assumes that each bound nucleotide is always in complex with a Mg^2+^ ion. A recent study, however, shows that the absence of Mg^2+^, especially in buffers with no EDTA, promotes KaiC autophosphorylation (Jeong et al., 2019). Moreover, some structures of KaiC have modeled two Mg^2+^ ions in the nucleotide-binding pocket, which has been interpreted to mean that KaiC kinase activity relies on a two-metal-ion phosphotransfer mechanism (Pattanayek et al., 2009). Currently, the functions of Mg^2+^ have not been characterized kinetically or mechanistically at a level necessary to constrain a molecularly detailed model such as ours.

### The hexameric structure of KaiC

The current model does not consider any hexameric effect. There is evidence to suggest that intersubunit interaction regulates KaiC autokinase activity (Kitayama et al., 2013) as well as the ultrasensitive dependence of KaiBC complex formation on the KaiC hexamer phosphoform composition (Lin et al., 2014). However, explicitly accounting for the hexameric nature of KaiC, as in Li et al. (2009) or Lin et al. (2014), would lead to a significant increase in the number of model parameters, which likely cannot be constrained by available data and makes interpretation of the model difficult. Therefore, we only keep track of monomeric KaiC species, and the rate constants should be considered averages over hexameric background configurations, weighted by their nonequilibrium state populations.

This leads to two simplifications concerning the KaiA-KaiC interactions. The first issue relates to the stoichiometry of KaiAC complexes. During phosphorylation, KaiA dimers bind to KaiC hexamers with either a 1:1 or 2:1 stoichiometry (Hayashi et al., 2004a; Yunoki et al., 2019). Because the model does not consider the hexameric structure of KaiC explicitly, this stoichiometry is not enforced, and each KaiC monomer can bind independently to KaiA.

The second issue relates to the effect of the hexameric phosphorylation state on KaiA (un)binding kinetics. KaiA and KaiB compete with each other for KaiC binding (Lin et al., 2014), even though their binding sites are on opposing sides of KaiC. This has been interpreted as a result of cooperative allosteric transition between a kinase mode of KaiC, stabilized by KaiA binding and the T phosphoform, and a phosphotase mode of KaiC, stabilized by KaiB binding and the S phosphoform (Lin et al., 2014). An implication of this proposed mechanism is that KaiC hexamers in the phosphatase mode have uniformly diminished affinity for KaiA at the CII interface, regardless of the phosphorylation state of the subunits. Given that the current model is trained using primarily the phosphorylation data set, the predicted KaiA dwell time (**Figure S4**B) and dissociation constant (**Figure 1**D) likely reflect the property of KaiC subunits in the kinase mode, whereas experiments with phosphomimetic mutants mimicking the S and D phosphorylation states presumably probe the system kinetics in the phosphatase mode. Indeed, in the KaiC EE titration experiment (**Figure 3**F right), the presence of KaiC EE has virtually no effect on the KaiC S431A stimulus-response curve, which may be partly due to the fact that the KaiC EE hexamers are in the phosphotase mode, a condition not considered in our model.

### Correlation structure in the posterior distribution

As discussed in the literature Gutenkunst et al. (2007), often ratios of parameters are better constrained than the parameters themselves. The parameter pairs that have a correlation coefficient larger than 0.9 in log space are shown in **Figure S5**A-C. Such correlations typically reflect that thermodynamic, rather than kinetic, properties of the system are constrained. These include the free energy of phosphotransfer between the S and D phosphoforms (**Figure S5**A) and the free energy for KaiA binding to the ATP-bound states of KaiC (**Figure S5**B; compare with **Figure 1**D). Interestingly, there is a linear relation among 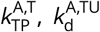, and 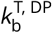 (**Figure S5**C); this implies that the data constrain the flux out of the 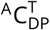 state, but the exact pathway is underdetermined.

More generally, we characterize the extent to which the model parameters, or linear combinations thereof, are constrained by the data using the principal components of the posterior distribution; that is, the eigenvectors of the covariance matrix from MCMC sampling. The eigenvalues of the covariance matrix span multiple orders of magnitude with no obvious gap (**Figure S5**D left), except for the stiffest direction (**Figure S5**G), which is almost entirely aligned with *σ*^2^, the global error hyperparameter. In addition, the directions of the principal components are in general not aligned with the directions of the bare coordinates (**Figure S5**D right), and there is no obvious interpretation for the directions of most of the principal components (**Figure S5**G). These features of the ensemble indicate that the model is “sloppy” (Gutenkunst et al., 2007), and many model parameters are poorly constrained by the data. Nevertheless, as we demonstrate in Results, the model can still be used to make consistent predictions because the variabilities in the ensemble of parameter sets obtained from MCMC sampling align with the softest degrees of freedom of the posterior distribution.

### Convergence of the model fit

We assessed the quality of the fit in three ways. First, we repeated the full procedure three times to assess reproducibility of the fitting procedure (**Figure S1**B). The three independent runs yielded marginalized posterior distributions that are remarkably consistent and tightly constrained for some parameters but diverge over several orders of magnitude for others. The best fits from the three runs have log posterior values of 720, 714, and 705, respectively. Unless otherwise specified, in this work we base our analyses on the run that produced the best fit with the highest posterior value. The ruggedness of the posterior distribution demonstrates that given the model and training data set, the parameter estimation problem is far from the asymptotic (i.e., large sample) regime. Moreover, in our fitting procedure the vast majority of the walkers from the annealing step is discarded for the sake of improving the MCMC sampling efficiency (the acceptance rate is ≤ 7% without pruning). Given the presence of multiple local maxima, this choice likely resulted in an underestimation of the uncertainties in parameter values.

**Table S1:** Effects of KaiA on KaiC function

| Model* | Log likelihood |  |  | Bayes factor |
| --- | --- | --- | --- | --- |
|  | Phosphorylation | Dephosphorylation | Hydrolysis |  |
| Full model | 422.9 | 346.8 | -0.8 | 1 |
| -P<sup>†</sup> | 390.0 | 338.6 | -0.8 | 1.4 |
| -H<sup>‡</sup> | 355.5 | 308.8 | -0.9 | 2.5 |
| -P,-H | 278.2 | 283.6 | -0.2 | 7.7 |

\* We do not consider a model where KaiA binding does not promote nucleotide exchange, because such models are incapable of autophosphorylation by construction.
<sup>†</sup> -P: KaiA binding decoupled from (de)phosphorylation rates.
<sup>‡</sup> -H: KaiA binding decoupled from KaiC hydrolysis rates.

Second, we compared the model predictions with a test data set that probed the phosphorylation reaction at two non-standard KaiC concentrations (**Figure S4**E). The fit quality on the test data set is somewhat worse compared to the training set (compare with **Figure 1**B). In particular, the model overestimates the D phosphoform concentration at 1.75 μM KaiC and underestimates the T and D phosphoform concentrations at 7 μM KaiC. This result suggests some degree of overfitting. This, however, is not a significant issue because we base our conclusions on the ensemble of walkers rather than the behavior of the best fit.

Lastly, we assessed the convergence of the MCMC simulations using the integrated autocorrelation times for the 48 principal components of the posterior distribution (**Figure S5**E). The autocorrelation times for the largest and smallest principal components are 8,500 and 3,300 steps, respectively, which gives rough estimates of the times between independent samples for the slowest and fastest degrees of freedom. These estimates are far shorter than the length of the final MCMC runs in the fitting procedure (100,000 steps). We also checked the autocorrelation time for the KaiA binding affinities (**Figure S5**F), which is within the bound given by the principal components.

### KaiA function cannot be solely explained in terms of nucleotide exchange

Because of the generality of the model, the function of KaiA is not restricted to that of a nucleotide-exchange factor. In particular, the phosphotransfer and ATP hydrolysis rates are allowed to depend on KaiA binding. There is some experimental evidence to support such effects—KaiA binding inhibits dephosphorylation (Xu et al., 2003) and the addition of KaiA increases the ATPase activity of KaiC (Terauchi et al., 2007; Murakami et al., 2008). However, the biochemical mechanisms underlying these effects are not clear from the experiments; for example, does KaiA increase KaiC ATPase activity by reconfiguring the KaiC active site, or does KaiA binding indirectly promote ATP hydrolysis by shifting the KaiC population towards phosphoforms that have high ATPase activity?

To test whether KaiA binding has a direct effect on KaiC catalytic activities, we construct simplified models where hydrolysis and/or phosphotransfer is independent of KaiA binding and compare the resulting models to the full model using the Bayes factor (**Table S1**). We find that decoupling either phosphotransfer (model –P) or hydrolysis (model –H) from KaiA binding decreases the evidence for the simplified models, but the effects are weak, especially in comparison to a model where both classes of reactions are decoupled from KaiA binding (model –P,–H). These results indicate that the function of KaiA cannot be solely explained by nucleotide exchange, but we cannot conclusively distinguish between models –P and –H.

### The experimental data admit two S phosphorylation pathways

We analyzed a random selection of 500 walkers to understand the implications of their variations for the mechanisms of ordered phosphorylation. To simplify the analysis, we converted each walker to two singlesite models in which either T431 or S432 was available for phosphorylation but not both. We asked how important each reaction rate constant is to the overall T and S phosphoform concentrations in the single-site models using the relative first-order sensitivities computed at the standard reaction condition (i.e., 100% ATP with 1.5 μM KaiA). We focus on the initial phosphorylation rates because the steady-state rates are determined by balances of many contributing processes, making them harder to interpret. The parameter sensitivities at *t* = 1 h are used as proxies for the sensitivities of the initial phosphorylation rates.

Because each single-site model has 18 parameters, there are 36 sensitivities for the two phosphoforms. To characterize this high-dimensional space, we used spectral clustering (**Figure S6**A). Overall, the parameter sensitivities are much more constrained by the data for the T phosphoform than the S phosphoform, which is unsurprising given the relatively low concentrations of the S phosphoform under all experimental conditions in the training dataset. The clustering furthermore indicates that there are two plausible kinetic ordering mechanisms, which differ primarily in terms of the phosphorylation pathways taken by the S phosphoform (**Figure S6**A and B). In both clusters, the U → T transition is most sensitive to 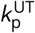 (i.e., the phosphorylation rate in the absence of KaiA).

In the first cluster (319 parameter sets), the U → S transition is most sensitive to 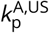 in the presence of KaiA. This is primarily because 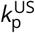 is very small in this cluster relative to 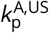 (**Figure S6**C); however, since, in this cluster, the *U* → *S* transition is dominated by the KaiA-bound states, the S phosphoform has negative sensitivity to the KaiA dissociation rate constant 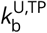. This suggests that in the first cluster, KaiA (un)binding to the U phosphoform is important in determining the initial phosphorylation rate of S. The best fit belongs to this first cluster.

The phosphorylation pathway suggested by the second cluster (181 parameter sets) is more complex. In this cluster, the U→S transition is mostly independent of KaiA, similar to the U→T reaction. However, the S phosphoform is limited by the dephosphorylation reaction 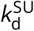, which is much faster than the corresponding phosphorylation rate (**Figure S6**C). In addition, the S phosphoform is sensitive to the rate constant for KaiA binding, 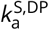, which is important for facilitating nucleotide exchange for the ADP-bound S phosphoform, but tends to be slower in the second cluster (**Figure S6**C). Therefore, faster dephosphorylation and slower KaiA binding is important for determining the initial S phosphorylation rate in the second cluster.

A comparison of **Figure S6**C with **Figure S1**B (blue distributions) shows that the two clusters correspond to the bimodal posterior distributions for the rate constants 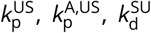, and 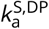. The two clusters, however, do not cleanly separate along the two modes of 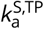 and 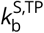; the kinetic significance of this bimodal distribution is unclear.

As discussed above, the posterior distribution is fairly rugged and thus the fitting procedure is not fully reproducible over independent runs. As a result, there are likely multiple potential kinetic ordering mechanisms that remain unexplored through this analysis. Regardless, the analysis suggests that kinetic ordering is likely a result of an interplay between (de)phosphorylation and KaiA (un)binding kinetics, rather than purely the product of equilibrium free energies of phosphotransfer.

### Comparison to the Paijmans model

Among all the computational work on the Kai oscillator, the model most similar to the current work in terms of the treatment of the KaiA-KaiC subsystem is that by Paijmans et al. (2017b), although the latter is a full oscillator model including KaiB, the CI domain, and the allosteric transition between the active (i.e., phopshorylation phase) and inactive (dephosphorylation phase) KaiC conformations. The Paijmans model and the full model in this work are both molecularly detailed, and describe how phosphotransfer, ATP hydrolysis, KaiA (un)binding, and nucleotide exchange reactions control the phosphoform and nucleotide-bound states of KaiC. However, there are some significant differences between these two models, which we examine below.

The Paijmans model is more general than this work in two ways. First, the Paijmans model explicitly considers the hexameric nature of KaiC. There is no intersubunit coordination of phosphotransfer in the Paijmans model, but it explicitly considers the binding of one KaiA dimer to a KaiC hexamer, which is assumed to uniformly accelerate nucleotide exchange in all six subunits. In this work, however, we do not consider the hexameric states of KaiC, and each KaiC monomer is allowed to bind to a KaiA dimer independently. In this way the affinities and kinetics of KaiA binding in this work may not be directly comparable to those in the Paijmans model. Second, the Paijmans model allows for the exchange of bound ATP for ADP, such that KaiA accelerates the exchange rates of both ATP and ADP while leaving the binding affinity unchanged. In our model, however, we assume that there is no exchange of bound ATP for ADP, effectively assuming that the affinity of ATP is infinite (i.e., 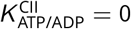 in the Paijmans model terminology). The treatment of nucleotide exchange in both models are otherwise similar, in that both assume that the ATP/ADP on rates are identical, that the apo state is in a quasi-steady state, and that there is no KaiA-independent nucleotide exchange.

The current work goes beyond the Paijmans model in the following ways. First, we determine the rate constants under the framework of Bayesian parameter estimation, which enables more rigorous uncertainty quantification, while the parameters in the Paijmans model were hand-tuned to reproduce selected experimental observations. Second, for simplicity the Paijmans model does not consider any possible coupling between ATP hydrolysis and KaiC phosphorylation states, between nucleotide exchange and KaiC phosphorylation states, between ATP hydrolysis and KaiA binding, or between KaiC nucleotide-bound states and KaiA binding. Although many of the species-dependent effects are not fully constrained by data in this work, as we describe in Results, the ultrasensitivity in KaiC phosphorylation depends critically on the coupling between KaiC nucleotide-bound states and KaiA binding. It is an open question whether a model that lacks such effects but explicitly accounts for the hexameric nature of KaiC can generate ultrasensitivity.

The difference in the treatment of KaiA binding affinity implies that some detailed balance conditions are incompatible between the two models. In the Paijmans model, the binding affinity of KaiA to KaiC hexamers during the phosphorylation phase depends on the phosphoform composition of the subunits, and each subunit *i* in the phosphoform X_*i*_ other than U contributes an additive factor of 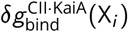 to the changes in KaiA binding free energy, 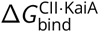. Due to detailed balance, the fact that KaiA binds to different KaiC phosphoforms with differential affinities implies that KaiA binding changes the free energy of phosphotransfer [see eq. 8 in Paijmans et al. (2017b)]. This condition is also present in our model, but is complicated by the nucleotide-bound states of KaiC. Using the multiplicative-factor parametrization scheme (see Materials and Methods), the detailed balance conditions in this work can be related to those in the Paijmans model by

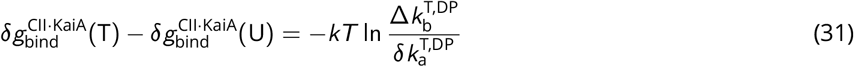

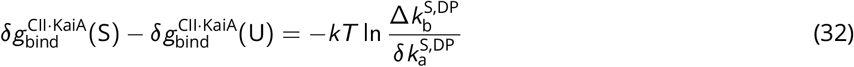

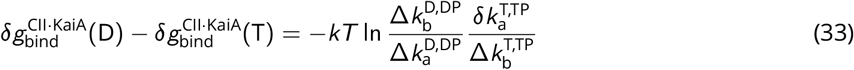

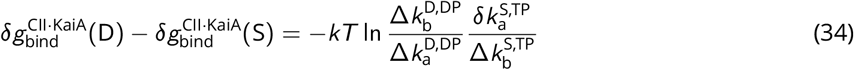

In general, this set of equations are inconsistent. That is, one cannot express the 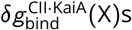 in the Paijmans model in terms of the Δ*k*s in our model. The only condition under which these equations can be made consistent is when

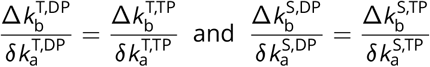

that is, when the nucleotide-bound states have no effect on KaiA binding affinities to the T and S phosphoforms.

### Phenomenological modifications to the Phong model

In the model by Phong et al. (2013), KaiA sequestration is determined by the equation

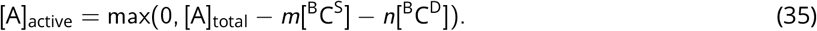

Here, [A]_total_ is the total KaiA concentration; [^B^C^S^] and [^B^C^D^] are the concentrations of the S and D phospho-forms in complex with KaiB, respectively; *m* and *n* are model parameters describing the binding stoichiometries between KaiA and the KaiBC complex. In this way, the KaiA binding affinity to the inhibitory complex is effectively infinite, with no KaiA dissociation until [^B^C^S^] and [^B^C^D^] drop below a threshold. To make the representation of KaiA sequestration more realistic, we introduce a KaiA dissociation constant *K*_*D*_ (**Figure 4**C). Assuming that the KaiA sequestration reaction is in a quasi-equilibrium, we replace (35) with

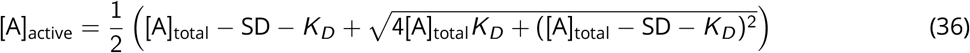

where we have defined SD = *m*[^B^C^S^] + *n*[^B^C^D^].

To introduce ultrasensitivity to the Phong model, we first note that the four phosphorylation rate constants for the U→T, U→S, T→D, and S→D transitions are given by Michaelis-Menten kinetics with ADP serving as a competitive inhibitor,

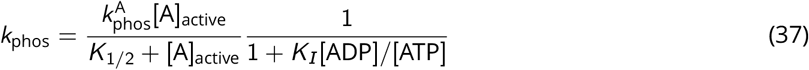

where 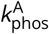 varies with the specific phosphorylation reaction. To introduce ultrasensitivity, we add a threshold term *T* (**Figure 4**D),

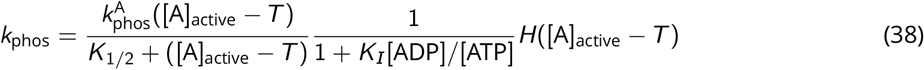

where

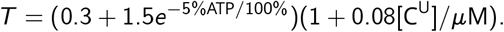

and *H* is the Heaviside function. The first part of the expression for the phosphorylation threshold, 0.3 + 1.5*e*^−5%ATP/100%^, describes how the threshold changes as a function of %ATP; the constants are determined by approximating the [KaiA] threshold in **Figure 3**A as an exponential function. The second part of the expression, 1 + 0.08[C^U^]/*μ*M, describes how the threshold changes as a function of C^U^ concentration. This formula is defined by

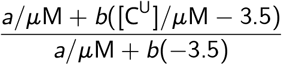

where parameters *a* and *b* are determined by taking a linear fit, *a* + *b*[AA], of the data from **Figure 3**G (yellow line); that is, 1 + 0.08[C^U^]/*μ*M gives the fractional changes to the phosphorylation threshold as a result of any U phosphoform (in the form of AA phosphomimetic mutant) additional to the 3.5 μM KaiC S431A in the experiment.

We use the original Phong model parameters in all analyses of the model with one exception. In the final model with both a *K*_*D*_ and a phosphorylation threshold (**Figure 4**E bottom), the period is systematically longer than 24 h due to a slow down in phosphorylation. To fix this problem, we change 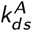 and 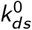, the two rate constants controlling the D→S transition, to 0.94 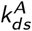 and 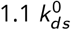.

---

**Figure S1:**
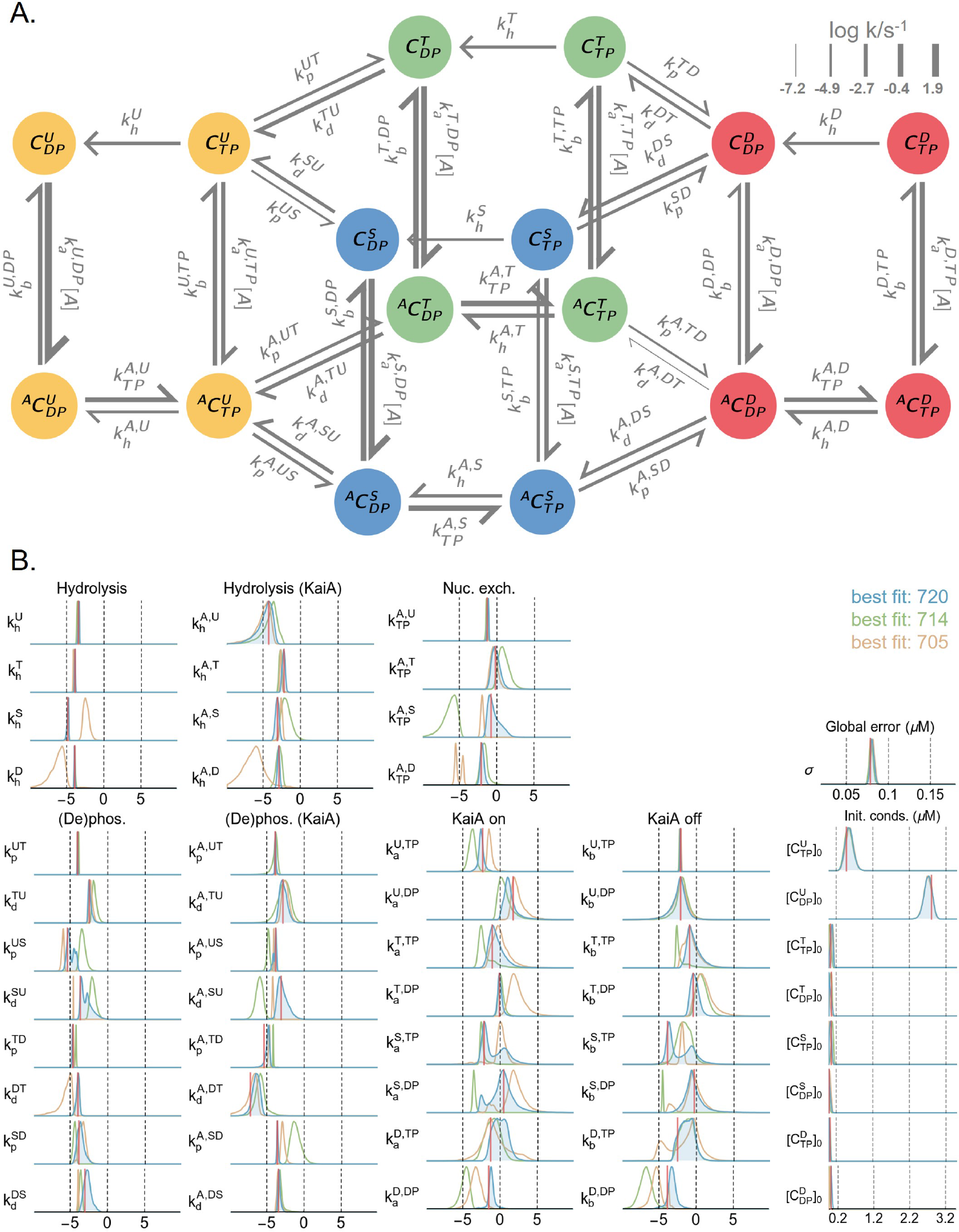
Overview of the model. A) A schematic of the full mass action kinetic model. Here, each arrow represents a reaction, and the associated rate constant is represented using the notation introduced in the main text. The thickness of the arrows is proportional to the best fit rate on a log scale (base 10) at 100% ATP and 1.5 μM KaiA. B) The posterior distributions for all rate constants, initial conditions, and the global error hyperparameter. The rate constants have the unit s^−1^(μM^−1^) and the horizontal axis has a log scale (base 10). The three distributions represent the results from three independent runs; the log posterior values for the best fits from the three runs are listed. The red lines represent the best fit from the best run (i.e., the blue distributions). See Materials and Methods and **Figure S10** for further details on the model parameterization method.

**Figure S2:**
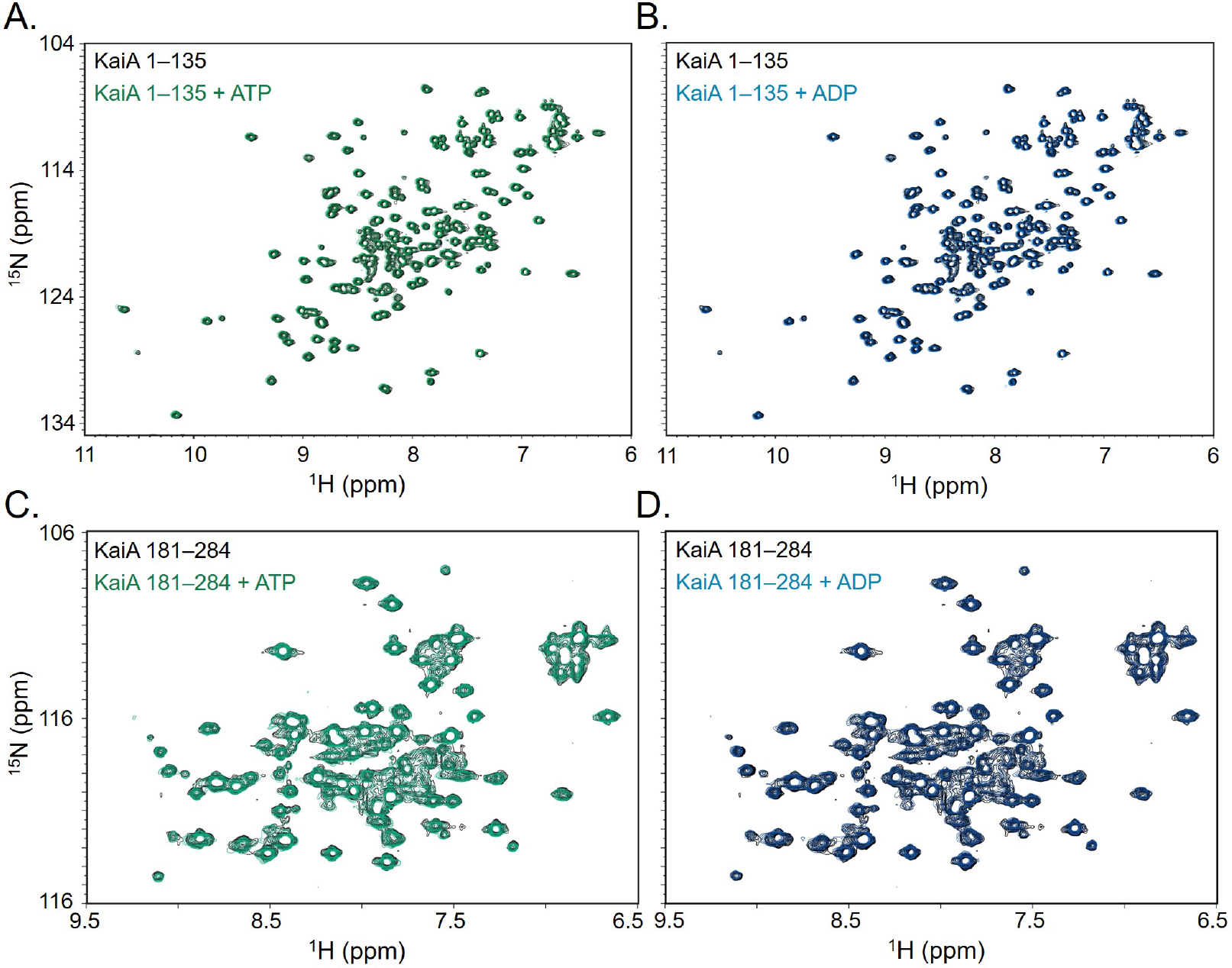
No evidence of direct nucleotide-KaiA interaction. ^1^H-^15^N HSQC spectroscopy of the N-terminal fragment (residues 1–135) of KaiA in the presence and absence of ATP (A) or ADP (B) show no significant differences in chemical shifts, while spectra of the C-terminal fragment (residues 181–284) show subtle line broadening in the presence of ATP (C) and ADP (D), suggesting weak, if any, interaction between the nucleotide and the C-terminal fragment. Given these results, we do not include any direct KaiA-nucleotide interaction in the model.

**Figure S3:**
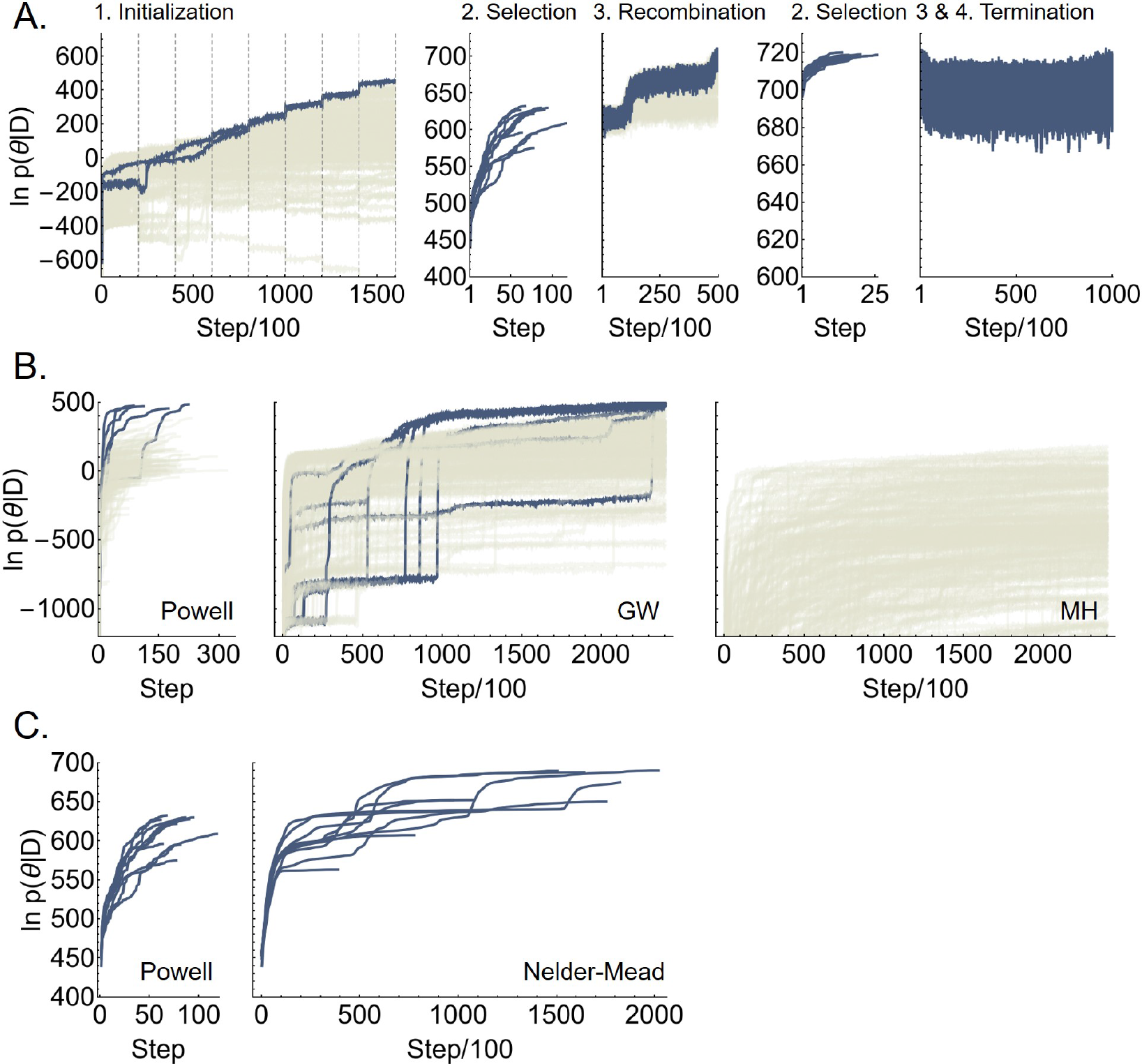
Performance of the fitting procedure. A) The time evolution of the log posterior values over the four steps of the fitting procedure (see Materials and Methods). For step 1 and 3, the individual Markov chains that do not produce walkers used in the next step are shown in beige. B) A comparison of the performance of Powell’s method, a derivative-free numerical optimization method, Goodman-Weare (GM), an ensemble MCMC method, and conventional MetropolisHastings (MH) algorithm with a Gaussian trial distribution. For the Metropolis-Hastings algorithm the covariance matrix of the trial distribution is given by the global covariance of the fit (i.e., the last step in panel A), scaled by a factor of 0.005 to give an average acceptance rate of 19.8%. A set of 224 walkers drawn from the prior distribution are used to initialize the simulations for all three methods; the 224 walkers are evolved independently for Powell’s method and Metropolis-Hastings, and in an ensemble for Goodman-Weare. Chains that do not reach log posterior above 450 are shown in beige. C) A comparison of the performance of Powell’s method with Nelder-Mead, a simplex-based numerical optimization method. The simulations are initialized using the same walkers as in step 2 of A).

**Figure S4:**
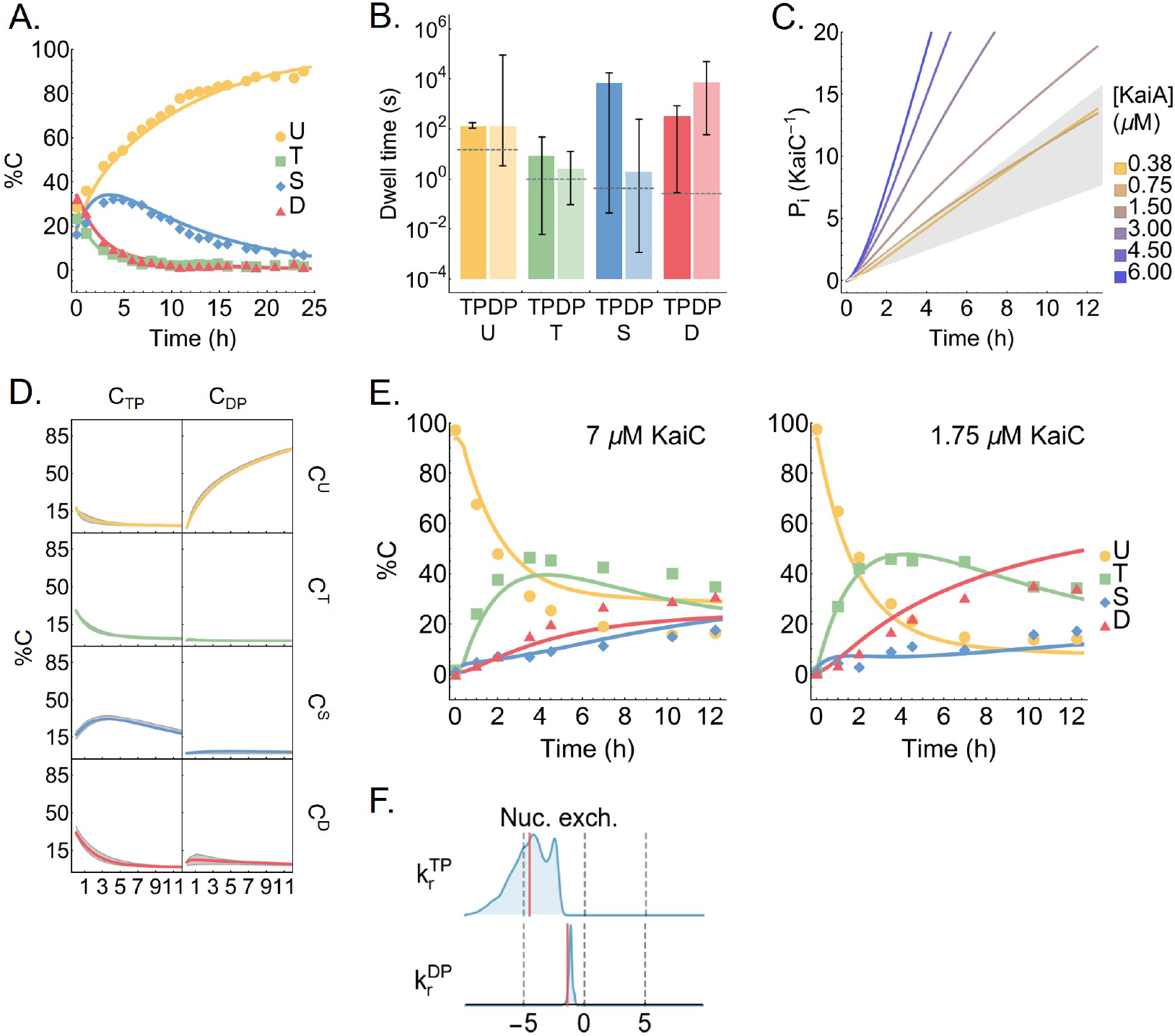
Behavior of the model. A) Model fit to the dephosphorylation dataset. B) The best fit KaiA dwell time as a function of KaiC phosphoform and nucleotide-bound state. The error bars represent the 95% posterior interval, and the dashed lines represent the experimental measurements, which did not resolve the nucleotide-bound states. C) Inorganic phosphate production per KaiC monomer over the course of a phosphorylation reaction. The gray region represents the experimental bounds on the KaiC hydrolysis rate with 1.2 μM KaiA and no KaiA. See Materials and Methods for the source of the experimental data in A–C. D) The kinetics of the dephosphorylation reaction in the absence of KaiA, broken down into the eight individual KaiC species. The gray region represents the 95% posterior interval. Refer to **Figure S1**A for the KaiC state names. E) The predicted phosphorylation kinetics at 7 and 1.75 μM KaiC, both at 100% ATP and 1.5 μM KaiA, compared to experimental measurements. Note that these two time series are not part of the training set. F) The posterior distributions for 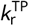 and 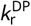, the dissociation rates for ATP and ADP, respectively, in an early iteration of the model. The rate constants have a unit of s^−1^ and the horizontal axis has a log scale (base 10). The long tail to the left of the posterior distribution for 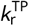 suggests that the model can be simplified by setting the rate to zero.

**Figure S5:**
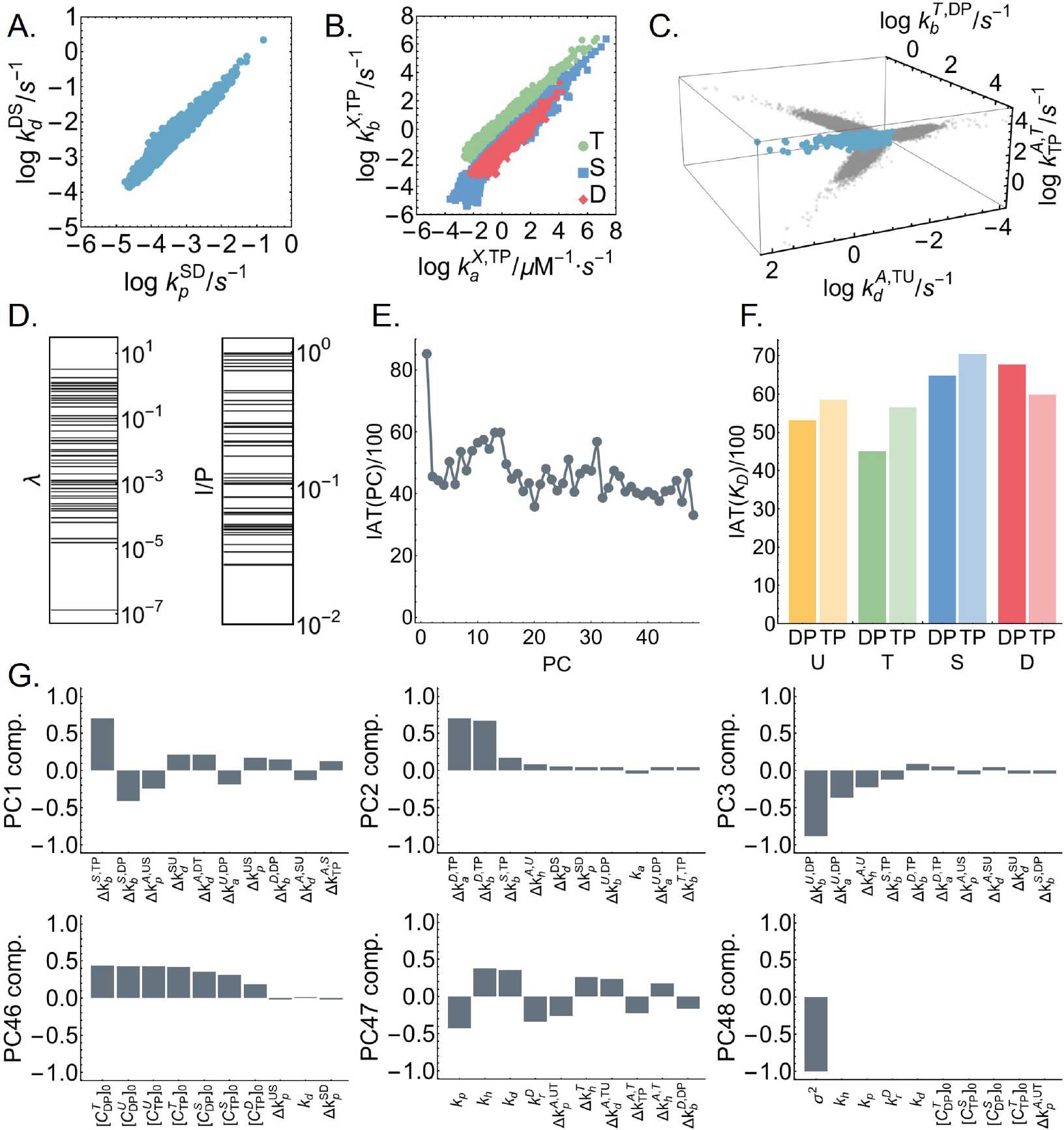
Correlation structure in the MCMC ensemble. A), B), and C) show parameters with a correlation coefficients larger than 0.9. In B), “X” represents the KaiC phosphoforms. In C), the projections of the 3D scatter plot onto pairwise correlations are shown in gray. D) The principal component/eigenvalue spectrum of the covariance matrix (left), and the alignment of the principal components with the coordinates (right). Here, *I* denotes the intersection of the principal component ellipoid with the coordinates and *P* denotes the projection of the principal components onto the corresponding coordinates (Gutenkunst et al., 2007). E) The integrated autocorrelation time for the 48 principal components (PC); the principal components are indexed from the largest to the smallest. The integrated autocorrelation time is calculated using an automated windowing procedure (Madras and Sokal, 1988) from the autocorrelation function averaged over the ensemble. F) The integrated autocorrelation time for the KaiA dissociation constants as a function of KaiC phosphoform and nucleotide-bound states. G) The ten largest vector components, ordered by absolute value, for the first and last three principal components.

**Figure S6:**
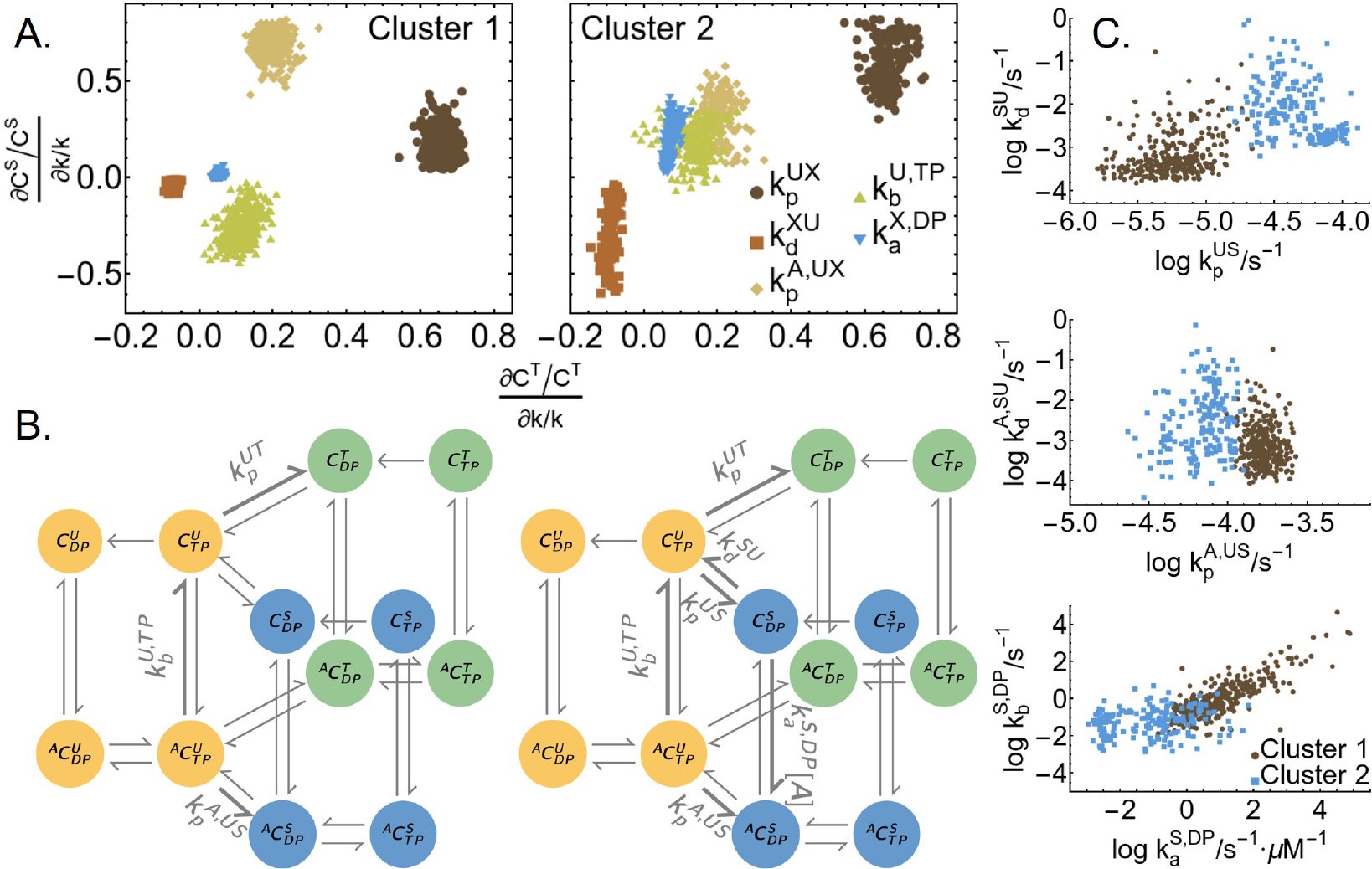
The mechanism of kinetic ordering is not well-constrained. A) Spectral clustering on the relative sensitivity of the T and S phosphoform concentrations at *t* = 1 h to rate constants in the T- and S-site models, respectively. Only the parameters with significant (> 0.2) relative sensitivities in either cluster are shown in the plot. “X” stands for either the T (horizontal axis) or S (vertical axis) phosphoform. The sensitivities are calculated using 500 sampled parameter sets chosen randomly from the ensemble. The clustering analysis was done using the FindClusters function in Mathematica 12.0. B) Model diagrams that highlight the reactions that have the highest relative sensitivities in the first (left) and second (right) clusters; the D phosphoform is not shown. C) Selected model parameter values in the two clusters. A comparison with blue distributions in **Figure S1**B indicates that the clustering based on sensitivity can be mapped onto the modes of the posterior distribution.

**Figure S7:**
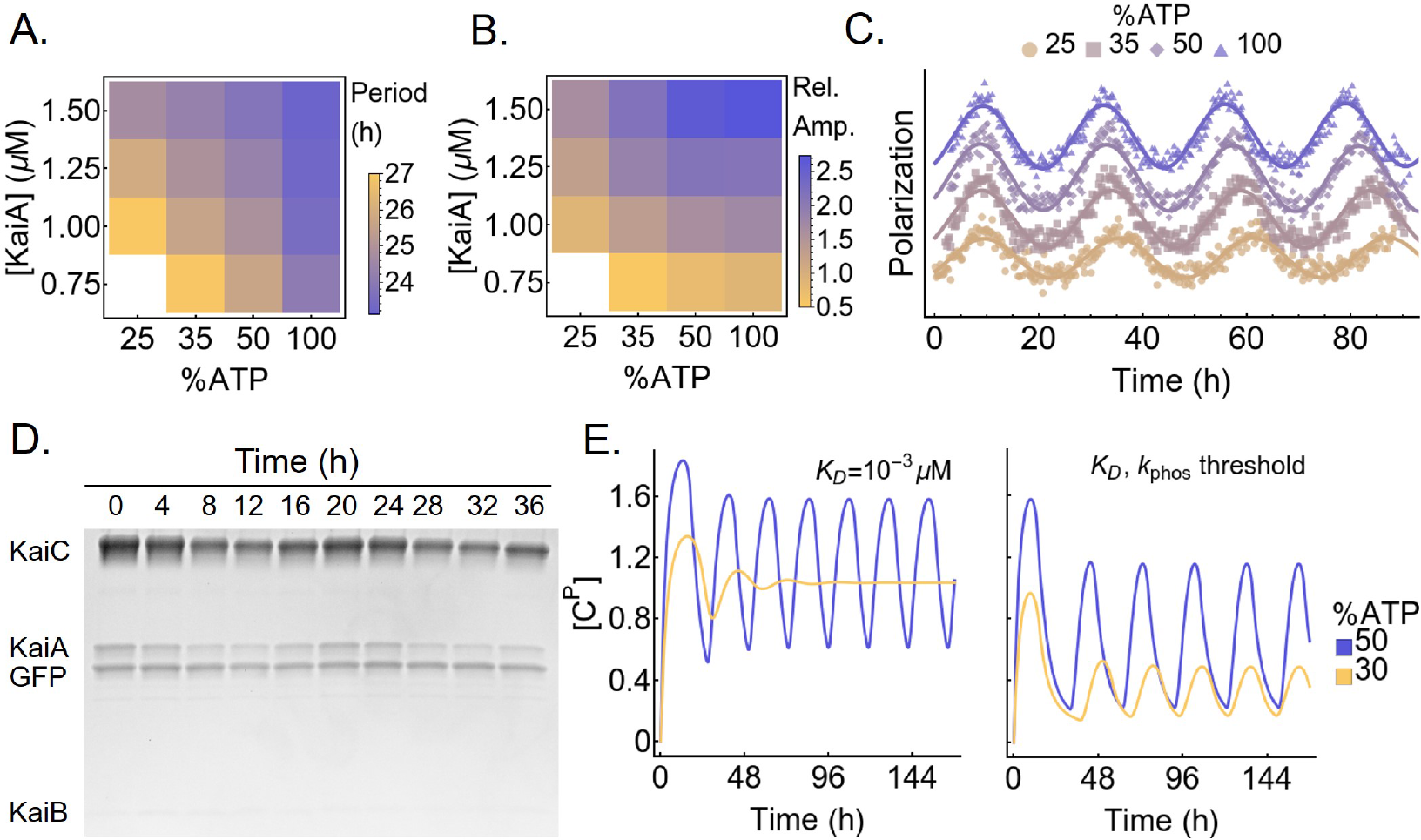
The metabolic compensation property of the Kai oscillator. The fluorescence polarization measurement of the oscillatory reactions are fit to a curve FP(*t*) = *A* cos(2*πT* ^−1^*t* + *ϕ*) + *bt* + *c* to extract A) the period (*T*) and B) the normalized amplitude (100*A/c*; dimensionless) of the oscillator as a function of [KaiA] and %ATP. Reactions with an amplitude *A* < 0.5 are considered to be non-oscillatory. C) Representative traces demonstrating the effect of %ATP at 1.25 μM KaiA; The polarization data are shifted vertically to avoid overlaps and horizontally to align the first peaks. D) SDS-PAGE gel image of the supernatant from the KaiB-FLAG immunoprecipitation experiment. E) A Comparison of the metabolic compensation property of the Phong model without (left) or with (right) a phosphorylation threshold at *K_D_* = 10^−3^ μM. The model exhibits phase decoherence at low %ATP without a phosphorylation threshold.

**Figure S8:**
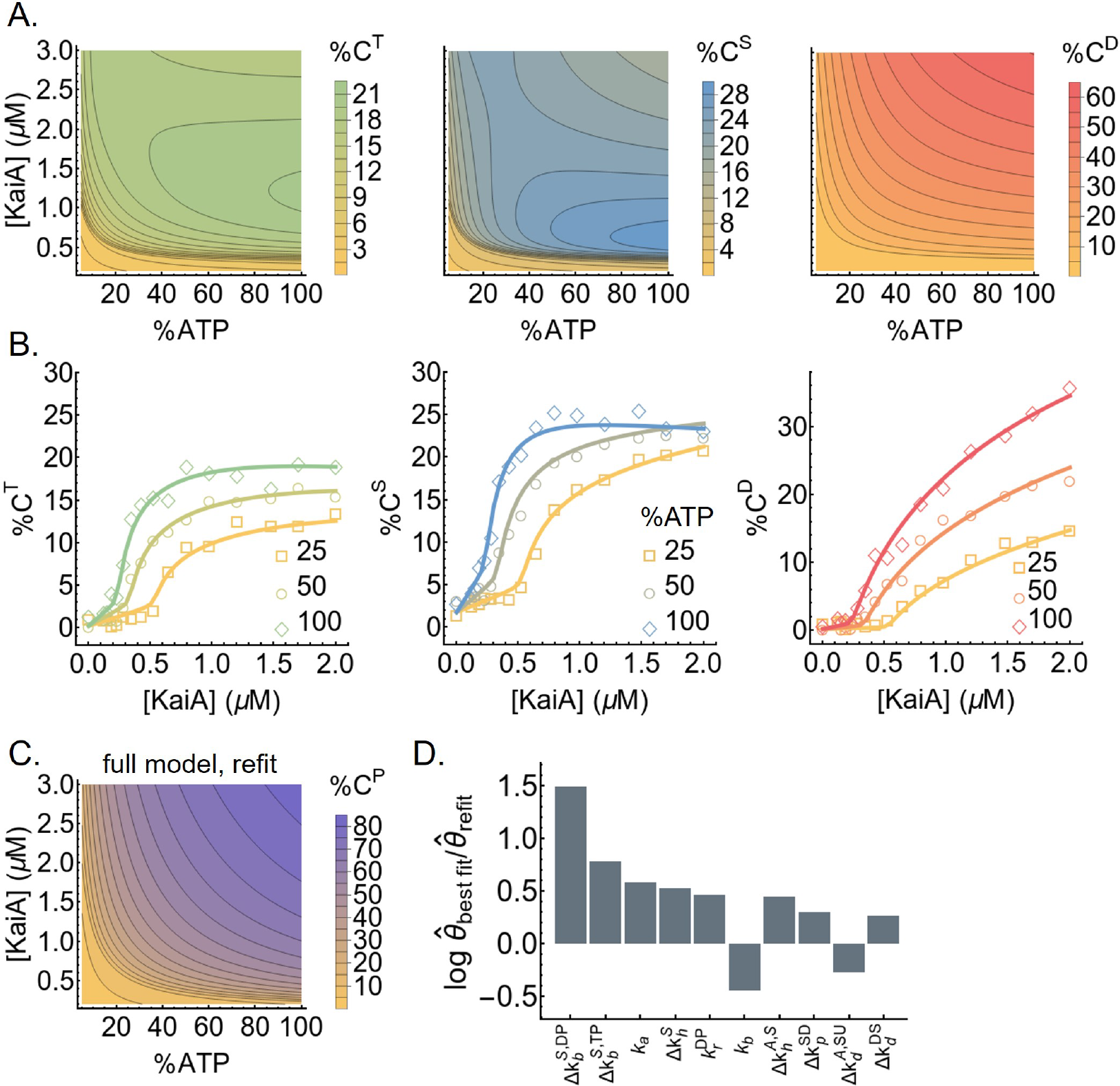
KaiC stimulus-response relations. A) The steady-state stimulus-response relations for T, S, and D phosphoforms predicted by the model. B) The experimentally determined stimulus-response functions of the T, S, and D phosphoforms at three %ATP conditions; the curves are based on refitting the best fit to the steady-state measurements. The model-predicted stimulus-response relation of the total steady-state KaiC phosphorylation level as a function of %ATP and [KaiA] after refitting to the steady-state measurements. D) The differences in the log parameter values (base 10) of the best fit before and after refit. The differences are ordered by magnitude and only the 10 parameters (in the multiplicative-factor scheme) with the largest changes are shown.

**Figure S9:**
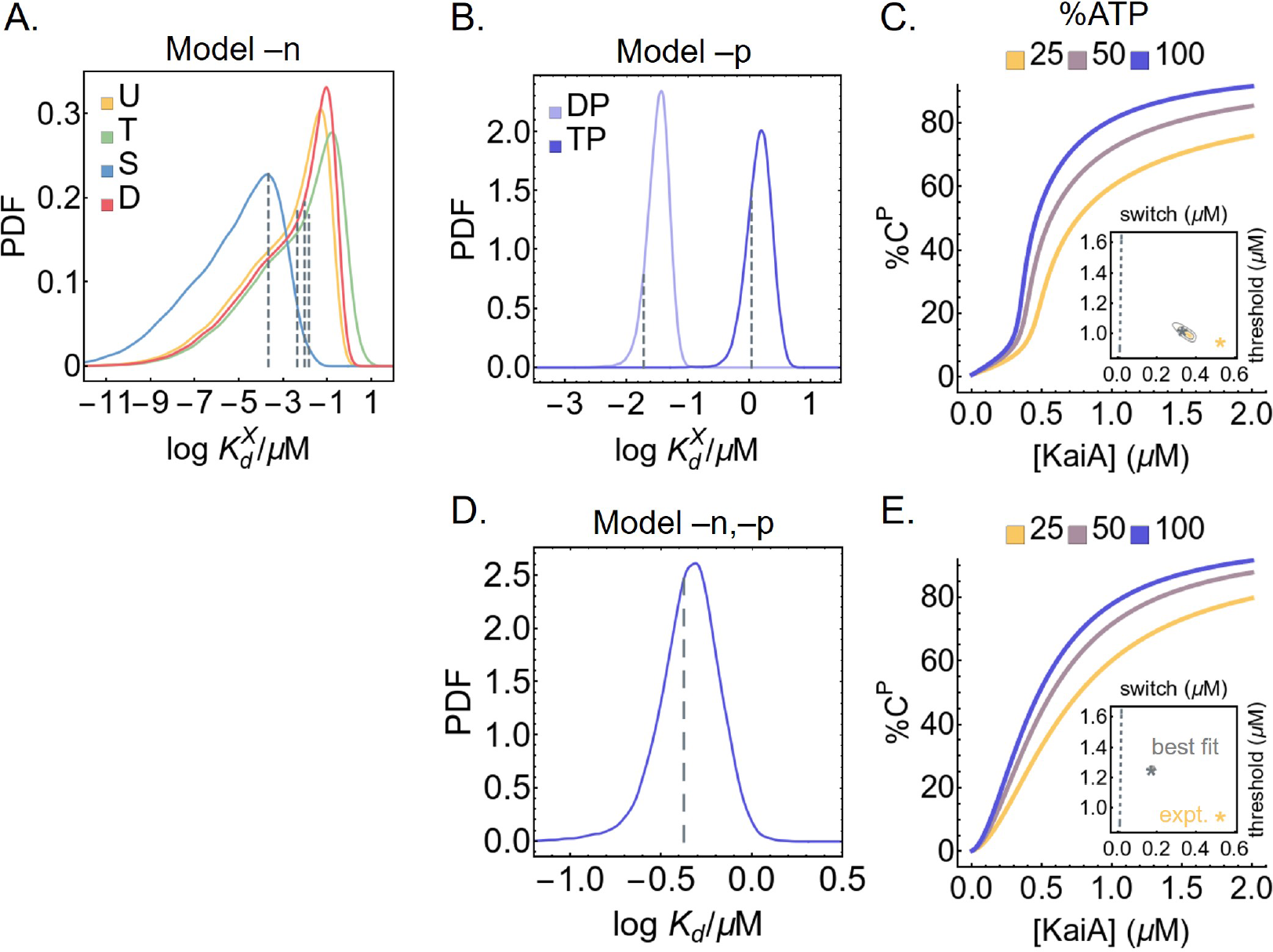
KaiA binding affinities of simplified models. A) The posterior distributions for the KaiA dissociation constants as a function of KaiC phosphoform in model –n, where the KaiA on/off rates are decoupled from the nucleotide-bound states of KaiC. The dashed lines represent the best fit. B) The posterior distributions for the KaiA dissociation constants as a function of KaiC phosphoform in model –p, where the KaiA on/off rates are decoupled from the KaiC phosphoform; the dashed lines represent the best fit. C) Cross sections of the stimulus-response relation at three %ATP, computed using model –p. The inset represents posterior distribution for the shapes of the stimulus-response function at 25% ATP. The contours represent the 68% and 95% HDRs, and the gray star represents the model best fit. The shape of the stimulus-response function is quantified using two metrics: EC10, which quantifies threshold-like behavior, and EC90 – EC10, which quantifies switch-like behavior. The shape of the experimentally-determined stimulus-response function at 25% ATP is shown as the yellow star. The dashed line represents (EC10, EC90 − EC10) = (*K*/9, 80*K*/9), which characterizes the shape of a hyperbolic stimulus-response function [A]/(*K* + [A]) that has no switching or thresholding. C) Similar to B), but for model –n,–p, where there is a single KaiA on/off rate in the model. E) Similar to C), but for model –n,–p.

**Figure S10:**
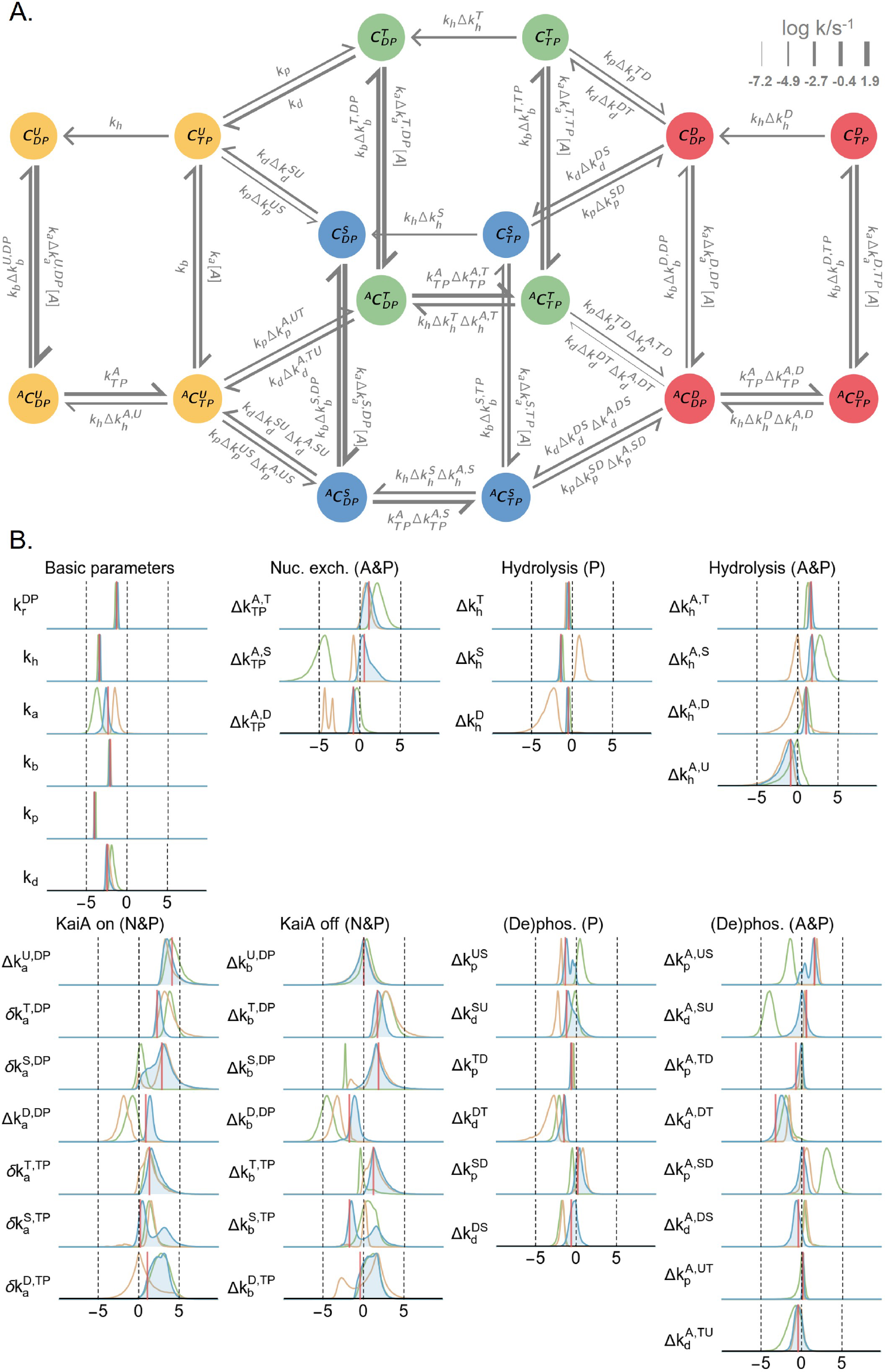
Overview of the model with the multiplicative-factor parameterization scheme. Panels A) and B) are analogous to those in **Figure S1**, but the rate constants are represented as products of the factors that are actually optimized in the MCMC simulations. In B), the *δk* parameters are fixed parameters determined by detailed balance conditions. The parentheses denote species-dependent effects; A: KaiA-bound state, P: phosphoform, N: nucleotide-bound state. See Materials and Methods for further description of the detailed balance conditions and the model parameterization method.

